# Molecular signature and functional properties of human pluripotent stem cell-derived brain pericytes

**DOI:** 10.1101/2023.06.26.546577

**Authors:** Ruslan Rust, Abhay P. Sagare, Kassandra Kisler, Youbin Kim, Mingzi Zhang, Casey Griffin, Yaoming Wang, Veronica Clementel, Arkadi Shwartz, Scott E Fraser, Carina Torres-Sepulveda, Gavin Spillard, Julia TCW, Berislav V. Zlokovic, Marcelo P. Coba

## Abstract

Brain pericytes maintain the blood-brain barrier (BBB), secrete neurotrophic factors and clear toxic proteins. Their loss in neurological disorders leads to BBB breakdown, neuronal dysfunction, and cognitive decline. Therefore, cell therapy to replace lost pericytes holds potential to restore impaired cerebrovascular and brain functions. However, the molecular composition and function of human iPSC-derived brain pericytes (iPSC-PC) remains poorly characterized. Here, we show by a quantitative analysis of 8,344 proteins and 20,572 phosphopeptides that iPSC-PC share 96% of total proteins and 98% of protein phosphorylation sites with primary human brain pericytes. This includes cell adhesion and tight junction proteins, transcription factors, and different protein kinase families of the human kinome. In pericyte-deficient mice, iPSC-PC home to host brain capillaries to form hybrid human-mouse microvessels with ligand-receptor associations. They repair BBB leaks and protect against neuron loss, which we show requires PDGRFB and pleiotrophin. They also clear Alzheimer’s amyloid-β and tau neurotoxins in an *ex vivo* brain slice assay via lipoprotein receptor. Thus, iPSC-PC may have potential as a replacement therapy for pericyte-deficient neurological disorders.

## Introduction

Pericytes are mural cells on brain capillaries that maintain blood-brain barrier (BBB) integrity and function ^1–5^. They contribute to formation of new blood vessels ^6^, secrete neurotrophic factors ^4^, and clear toxic proteins such as Alzheimer’s disease (AD) amyloid-β (Aβ) ^7^ and tau ^8^. Their loss in AD and related neurodegenerative disorders leads to BBB breakdown ^6^, is associated with synaptic and neuronal dysfunction ^3,4,9^ and can predict cognitive decline in individuals carrying the **ε*4* variant of apolipoprotein E (*APOE4*), the main susceptibility gene for AD ^10^. Therefore, replacing lost pericytes by induced pluripotent stem cell (iPSC)-derived therapies holds potential to restore impaired cerebrovascular and neuronal functions.

Different protocols have been developed to generate human iPSC-derived brain pericytes (iPSC-PC) ^11–14^. iPSC-PC have been shown to enhance BBB integrity in human *in vitro* BBB models ^11–13^ and in a mouse model of ischemic stroke ^13^. Additionally, it has been shown that iPSC-PC from APOE **ε*4/*ε*4* donors compared to **ε*3/*ε*3* donors lead to a greater Aβ deposition in human BBB models ^14^ and activation of BBB-degrading cyclophilin A-matrix metalloproteinase-9 pathway ^10^.

However, the molecular profile and functional properties of iPSC-PC are still poorly understood. Whether their molecular signaling machinery corresponds to primary adult human brain microvascular pericytes (PC) remains elusive. It is also unknown if the principal functional components of iPSC-PC including cell adhesion molecules, tight junction proteins, transcription factors, and the protein kinase families of the human kinome can recapitulate the signaling signatures and functions observed in PC. Moreover, in terms of function, whether iPSC-PC can home to brain capillaries that lost some of their own pericyte coverage to form new hybrid microvessels with the host endothelium, whether they can repair BBB leaks, and whether they can protect against neuronal dysfunction associated with pericyte deficiency, remains elusive, and the underlying molecular mechanisms remain unstudied.

To address these questions, we compared the molecular signatures of iPSC-PC and PC using a quantitative analysis of total proteome and phosphoproteome using liquid chromatography mass spectrometry (LC/MS) and bioinformatics. Next, we studied vasculogenic potential of iPSC-PC to form hybrid microvessels with the host brain capillaries in the hippocampus of pericyte-deficient mice, determined BBB integrity of newly-formed hybrid vessels, studied their effects on neuronal function, and evaluated their capability to remove Alzheimer’s Aβ and tau neurotoxins.

## Results

During *in vivo* development, CNS pericytes originate from either neuroectoderm or mesoderm. Neuroectoderm-derived neural crest cells (NCC) differentiate into pericytes of the forebrain while pericytes of the midbrain, brain stem, and spinal cord have mesodermal origin ^15^. Therefore, to generate human forebrain pericytes (iPSC-PC) from iPSC, we used a modified two-step approach with a NCC intermediate similar to previous reports ^11–13^ (**Fig. 1a**).

**Figure 1:**
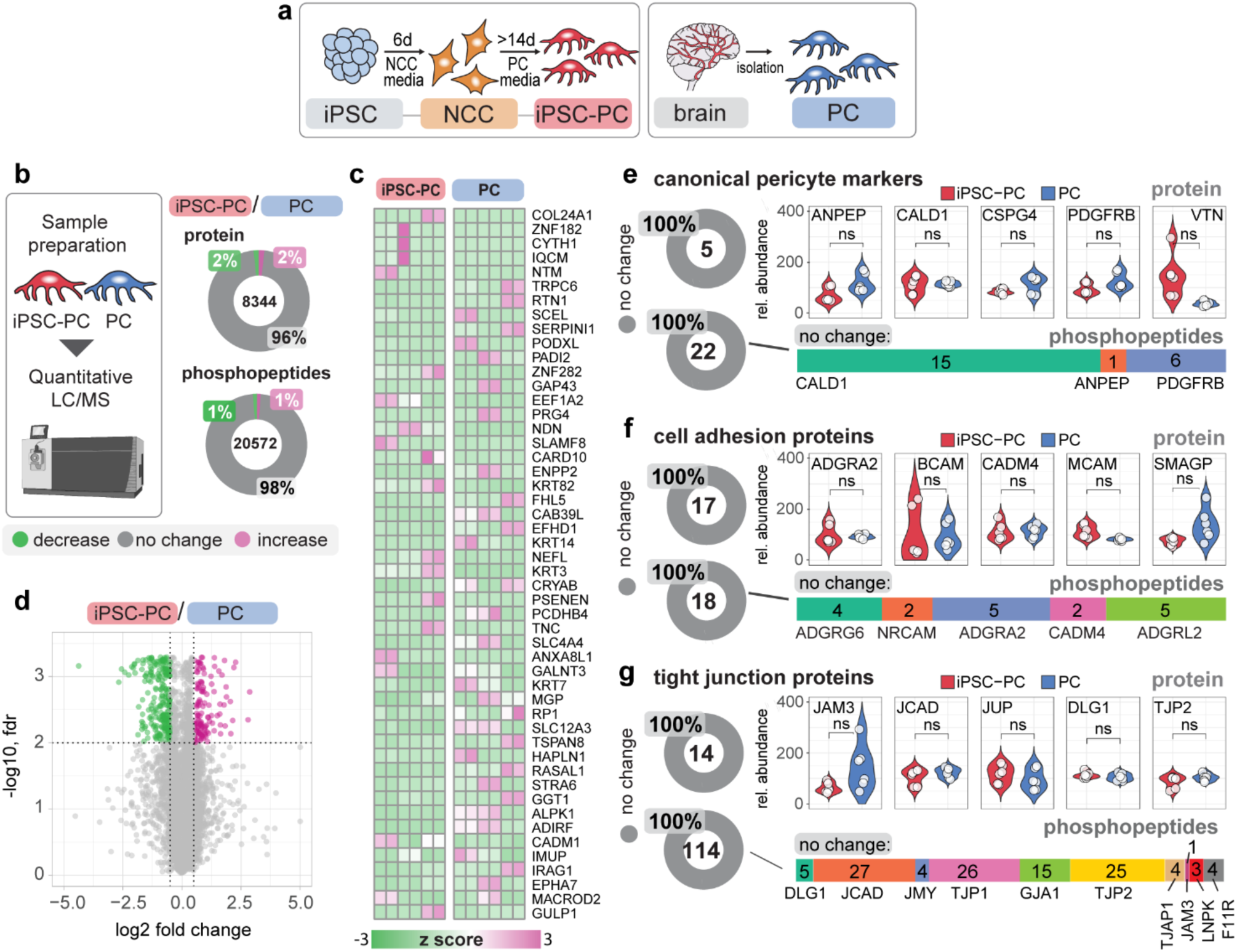
Generation and molecular characterization of iPSC-PC. (**a**) Schematic illustrating iPSC-PC were generated via a neural crest cell (NCC) intermediate, and primary human brain pericytes (PC) were isolated from postmortem cortical tissue. (**b**) Left: schematic illustrating proteomic data generation workflow by Liquid Chromatography/Mass Spectrometry (LC/MS). Right: Quantitative total proteome (top) and phosphoproteome (bottom) analysis of iPSC-PC vs. PC shown as donut plots (values inside plots represent total proteins/phosphopeptides; percentage indicates unchanged (grey), downregulated (green) and upregulated (magenta) proteins/phosphopeptides). Full lists of all identified proteins and phosphopeptides with relative abundance and q values can be found in Supp. Tables 1 and 2. (**c**) Heatmap of top 50 highly variant proteins between six independent cultures of iPSC-PC and PC. Protein expression is shown as a z score. Magenta indicates high expression, green indicates low expression. (**d**) Volcano plot of common and differentially expressed proteins identified between iPSC-PC/PC. Magenta dots indicate upregulation, grey dots indicate no change and green dots indicate downregulation. (**e-g**) LC/MS quantitative analysis of protein levels shown as donut plots and violin plots of individual biological replicates (top) and phosphorylation (bottom) of canonical pericyte markers (**e**), cell adhesion proteins (**f**) and tight junction proteins (**g**) in iPSC-PC (red) vs. PC (blue). In e-g, n = 6 independent cultures per group; Significance determined by two-stage Benjamini Yekutieli Krieger step up with multiple hypothesis. In **f** and **g**, 5 representative proteins are shown in violin plots. Full lists of protein and phosphopeptides for **e-g** can be found in Supp. Tables 3 and 4.

Immunostaining revealed that iPSC-derived NCC express NCC markers p75 (NGFR) and SOX10, but have barely detectable or undetectable levels of pericyte markers PDGFRB and NG2 (**Supp. Fig. 1a-c**). After differentiation into iPSC-PC, a robust expression of PDGFRB and NG2 was observed (**Supp. Fig. 1b-c**). Immunoblotting confirmed that iPSC-PC and PC express high levels of several canonical pericyte markers including NG2 (CSPG4), PDGFRβ (PDGFRB), CD13 (ANPEP), Caldesmon (CALD1), and Vitronectin (VTN) (**Supp. Fig. 1d**). As anticipated, markers of endothelial cells (GLUT1, CD31), neurons (NFL, TUJ1), oligodendrocytes (CNPase, OLIG2), astrocytes (ALDH1, GFAP, S100B), microglia (CD11B, CD68, IBA1) and epithelial cells (CDH1, KRT-18, EpCAM) were not detected in immunoblots of iPSC-PC or PC, when compared to their correspondent phenotypic cell-type controls (**Supp. Fig. 2a-g**). Only a faint signal of vascular smooth muscle cell markers (MYH11, ACTA2, CNN1) was observed in iPSC-PC and PC (**Supp. Fig. 2f**).

**Figure 2:**
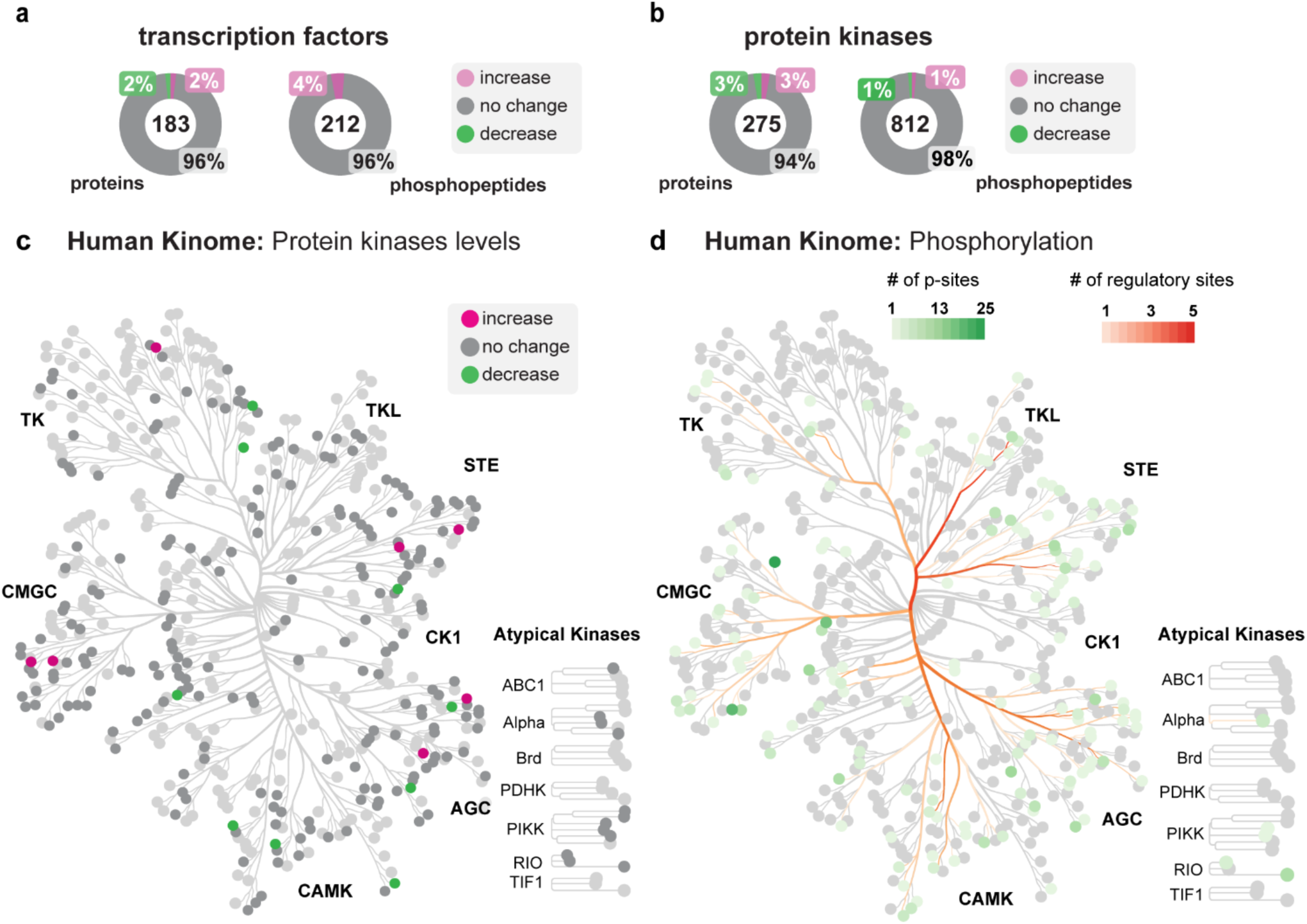
Proteomic analysis of transcription factors and protein kinases in iPSC-PC and PC. (**a**) Quantitative analysis of transcription factor protein levels (left) and phosphorylation (right) in iPSC-PC vs. PC by Liquid Chromatography/Mass Spectrometry (LC/MS). (**b**) LC/MS quantitative analysis of kinase protein levels (left) and phosphorylation (right) in iPSC-PC vs. PC. In **a**, **b**, data shown as donut plots (values inside plots represent total proteins/phosphopeptides; percentage indicates unchanged (grey), downregulated (green) and upregulated (magenta) proteins/phosphopeptides). Full lists of all identified proteins and phosphopeptides with relative abundance and q values can be found in Supp. Tables 1 and 2. (**c**) Distribution of protein kinases identified and quantitated in iPSC-PC vs. PC within the human Kinome. Proteins with no changes (gray) and changes (magenta or green) in total protein levels are shown. Light gray: not detected. (**d**) Distribution of unchanged protein kinase phosphorylation sites in iPSC-PC and PC. Green nodes: number of phosphorylation sites identified per protein kinase. Red branches: number of known regulatory sites per individual protein kinase node at the termination of the branch. Light gray: not detected. In **a-d**, n = 6 independent cultures per group; Significance determined by two-stage Benjamini Yekutieli Krieger step up with multiple hypothesis indicated. See Supp. Tables 3 and 4 for details. In **c** and **d**: TK: Tyrosine kinase group; TKL: Tyrosine kinase-like group; STE: Sterile (homologs of yeast Ste7) group; CK1: Casein Kinase 1 group; AGC, Protein kinase A, Protein kinase B, Protein kinase C group; CAMK: Calmodulin calcium kinase group; CMGC: [cyclin-dependent kinases (CDKs), mitogen-activated protein kinases (MAPKs), glycogen synthesis kinases (GSKs), and CDK-like kinases] group. Full lists of transcription factor and protein kinase proteins, phosphopeptides, and kinase phosphosites can be found in Supp. Tables 3 and 4.

LC/MS analysis performed as we previously reported ^9,16–19^ allowed us to identify and quantitate 8,344 proteins and 20,572 phosphopeptides showing no significant changes between iPSC-PC and PC (q>0.01) in 8,027 proteins (96%) and 20,135 phosphopeptides (98%) (**Fig. 1b-d, Supp. Tables 1, 2**). All proteomic and phosphoproteomic data are accessible via ProteomeXchange data base. Heatmap analysis of the top 50 most variably expressed proteins revealed a homogenous protein expression between iPSC-PC and PC, lacking any distinct pattern segregating the two groups (**Fig. 1c**). Volcano plot of common (8,027) and differentially expressed upregulated (134) or downregulated (183) proteins between iPSC-PC vs PC is shown in **Fig. 1d**. Full list of all identified and quantitated proteins and phosphopeptides and q values can be found in **Supp. Tables 1** and **2**.

LC/MS quantitative analysis of canonical pericyte markers including PDGFRβ (PDGFRB), Caldesmon (CALD1), NG2 (CSPG4), CD13 (ANPEP), and Vitronectin (VTN) indicated no changes in total protein levels or within 22 phosphopeptides between iPSC-PC and PC (**Fig. 1e, Supp. Tables 3, 4**). Importantly, we did not observe changes in phosphorylation in regulatory sites on PDGFRB S712 and S1104, which are essential for PDGFRB activity and association in protein interaction networks.

We then asked if the same pattern is observed in the main functional components of iPSC-PC. Therefore, we analyzed changes in the total protein and protein phosphorylation in cell adhesion and tight junction proteins, transcription factors, and protein kinases. We observed no changes in total protein and phosphorylation levels within 17 cell adhesion proteins and 18 phosphopeptides (**Fig. 1f, Supp. Tables 3, 4**). The same pattern was observed for tight junction proteins with no changes in protein levels in all studied proteins and all 114 phosphopeptides detected (**Fig. 1g, Supp. Tables 3, 4),** including major components of the pericyte tight junctions such as TJP2, TJP3, GJA1, DLG1 and others. These results suggest that cell adhesion and tight junction functions are highly preserved in iPSC-PC.

To determine if the intracellular signaling machinery and output response of iPSC-PC resemble that of PC, we analyzed total protein and phosphorylation changes in the transcription factors and protein kinases detected. We observed no significant changes (q>0.01) in total protein content in 96% of 183 total transcription factors, nor in protein phosphorylation in 96% of 212 identified phosphopeptides within 52 transcription factors (**Fig. 2a, Supp. Tables 3, 4**), suggesting no changes in the output response regulating total protein content in iPSC-PC. The analysis of the pericyte kinome allowed us to identify and quantitate a total of 275 protein kinases among all branches of the human kinome, including TK: Tyrosine kinase group; TKL: Tyrosine kinase-like group; STE: Sterile (homologs of yeast Ste7) group; CK1: Casein Kinase 1 group; AGC: Protein kinase A, Protein kinase B, Protein kinase C group; CAMK: Calmodulin calcium kinase group; CMGC [cyclin-dependent kinases (CDKs), mitogen-activated protein kinases (MAPKs), glycogen synthesis kinases (GSKs), and CDK-like kinases] group. We observed no significant changes (q>0.01) in 94% of protein kinases (**Fig. 2b-c, Supp. Tables 3, 4**). We also identified and quantitated 812 phosphopeptides with 636 unique phosphorylation sites within 160 protein kinases distributed among different families of the human kinome (**Fig. 2b,d**, **Supp. Tables 3, 4**) with no significant changes (q>0.01) in 98% of phosphopeptides.

The activity of protein kinases is usually modulated by protein phosphorylation of key phosphorylation sites that regulate their intrinsic kinase activity, protein localization and association to protein partners ^20^. Therefore, we identified protein kinase regulatory sites within the protein phosphorylation sites quantitated using the PhoshoSitePlus database resource followed by manual curation ^21^. We were able to identify 117 regulatory phosphorylation sites distributed along different protein kinase families in the pericyte kinome. Protein quantitation showed no significant differences (q>0.01) in any of the identified regulatory phosphorylation sites (**Fig. 2d, Supp. Tables 3, 4**). Among the 317 proteins and 437 phosphopeptides that were differentially expressed between iPSC-PCs and primary PCs, the changes appeared functionally diverse and did not converge on a specific pathway (**Supp. Tables 3, 4).** Altogether, these findings provide strong support for the high degree of similarity in the molecular signatures and functional capacity of iPSC-PC and PC, and suggest that iPSC-PC not only express similar protein levels in the main functional components of their total proteome as PC, but their protein functional characteristics are also preserved as we did not observe major changes in their phosphorylation signatures.

We next performed *in vitro* and *in vivo* experiments with iPSC-PC to evaluate their physiological functions. Since brain pericytes are known to enhance tight junction (TJ) formation between brain endothelial cells ^2,3^, we cultured primary human brain endothelial cell (hBEC) monolayers under three conditions: with iPSC-PC conditioned media, PC conditioned media, or unconditioned media for 24 h (**Fig 3a,b**). We then stained hBEC monolayers for TJ proteins claudin-5 (CLDN5) and ZO-1 (TJP1). Consistent with previous reports showing that iPSC-PC can enhance integrity of endothelial monolayers in human *in vitro* BBB models ^11–13^, claudin-5 (**Fig. 3c-d**) and ZO-1 (**Fig 3e-f**) had significantly longer TJ length (p < 0.05) in EC monolayers exposed to iPSC-PC or PC conditioned medium, compared to BEC medium only. To confirm the hBECs did not undergo a phenotypic shift in culture, we also verified that hBECs lacked epithelial markers (**Supp. Fig. 3**). Notably, similar effects on TJ proteins were also observed with cultured mouse brain endothelial cells (mBEC) monolayers conditioned with iPSC-PC media (**Supp. Fig. 4).** To further demonstrate that iPSC-PC exhibit functional properties similar to PC, we assessed their physiological response to KCl treatment, which triggers cell contraction through membrane depolarization ^22^ (**Fig. 3g**). Both iPSC-PC and PC showed a 30% change in length 20 minutes after KCl treatment compared to untreated controls (**Fig. 3h-i**), indicating a similar contractile response.

**Figure 3.**
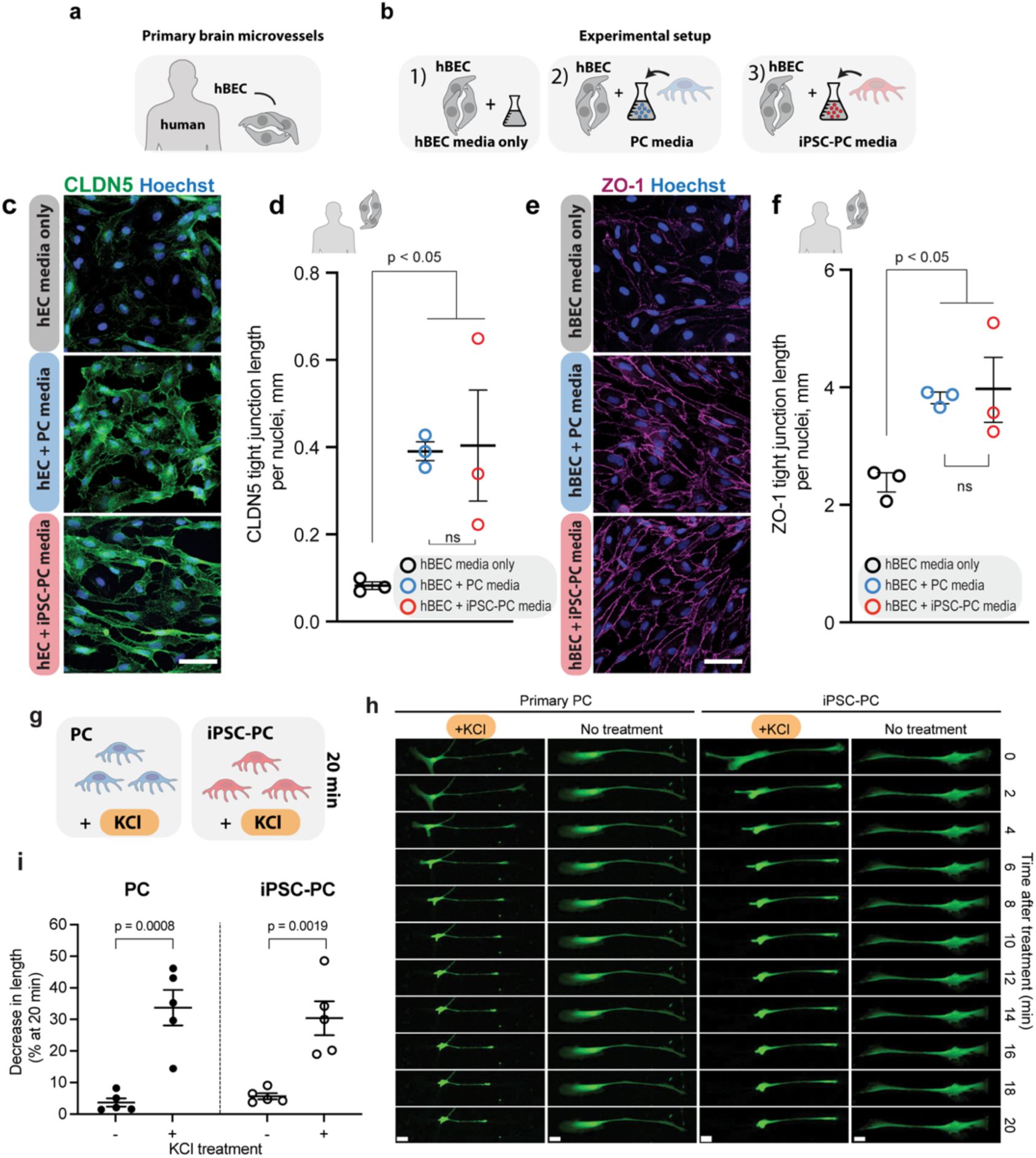
Impact of iPSC-PC on blood-brain barrier tight junction protein length and contractility. (**a,b**) Schematic overview of experimental design using primary human brain endothelial cells (hBEC) (**a**) with different culture media conditions (**b**). (**c,d**) Representative images of tight junction protein Claudin-5 staining in human brain endothelial cell (hBEC) cultures after 24 h treatment with iPSC-PC conditioned media, PC conditioned media or hBEC media only (**c**), and quantification of Claudin-5 length normalized to total number of Hoechst nuclei (**d**). (**e,f**) Representative images of tight junction protein ZO-1 staining in hBEC cultures after 24 h treatment with iPSC-PC conditioned media, PC conditioned media or hBEC media only (**e**), and quantification of ZO-1 length normalized to total number of Hoechst nuclei (**f**). (**g-i**) Schematic showing physiological depolarization of iPSC-PC using 20 min KCL treatment of iPSC-PC and PC (**g**), representative time-lapse images of PC and iPSC-PC cell contraction in response to KCl vs untreated controls (**h**), and quantification showing changes in length of PC and iPSC-PC following KCl treatment (**i**). In **d**, **f**, **i** data presented as mean ± SEM; n ≥ 3 replicates, with replicates represented by circles. Significance by Dunnett’s multiple comparisons test (d,f) and unpaired t test (i). Scale bars: (**c, e**) 25 µm, (**h**) 20 µm.

**Figure 4.**
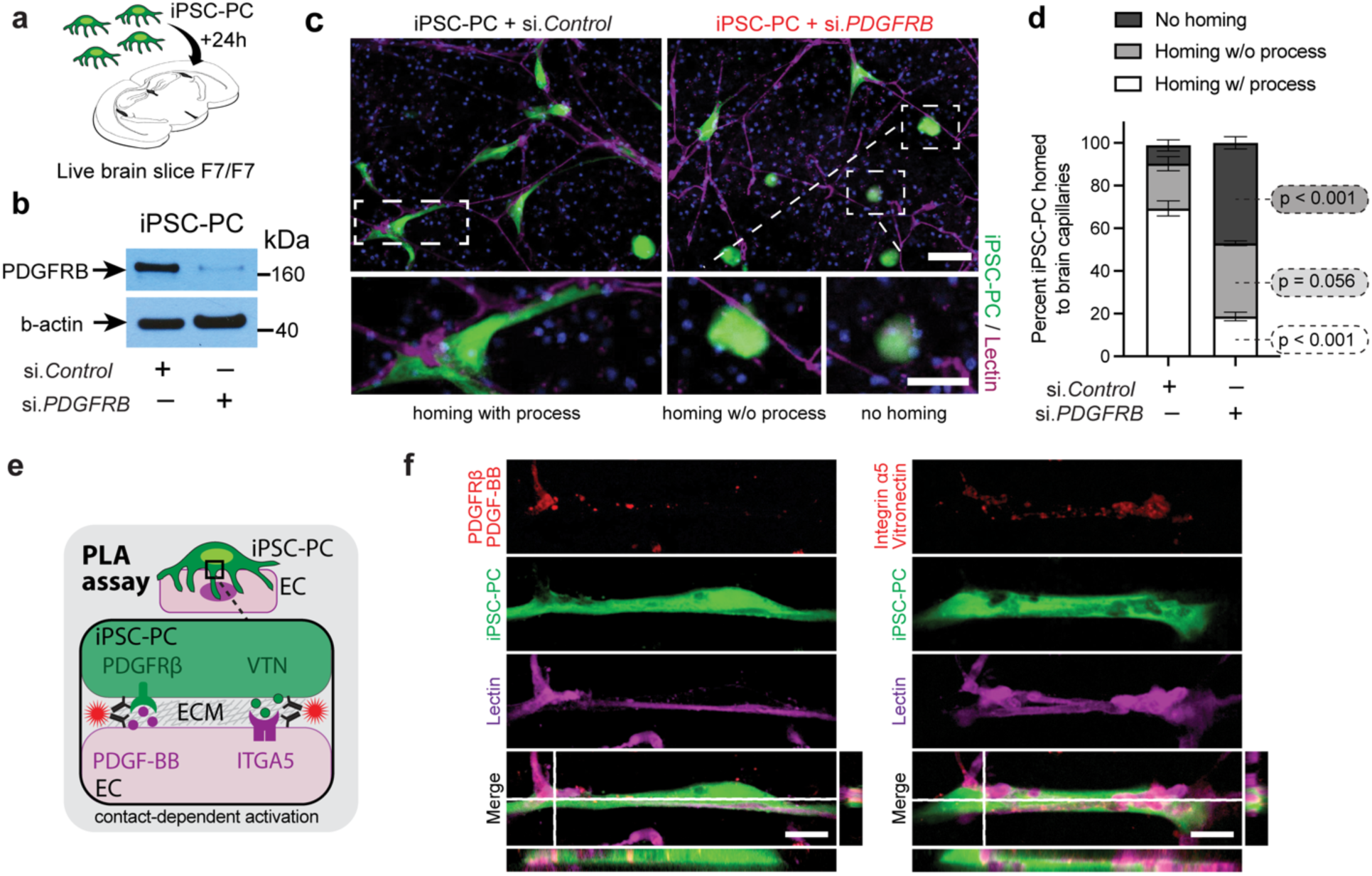
iPSC-PC home to brain capillaries of pericyte-deficient mice to form hybrid human-mouse brain capillaries. (**a**) Schematic showing the *ex vivo* experimental model. CellTracker-labeled iPSC-PC were incubated with live brain slices from pericyte-deficient *Pdgfrb^F^*^7^*^/F^*^7^ (F7/F7) mice for 24 h. (**b**) Immunoblotting for PDGFRB in iPSC-PC transduced with either control siRNA (si.*Control*) or human *PDGFRB* siRNA (si.*PDGFRB*). (**c-d**). Integration of CellTracker-labeled iPSC-PC (green) transduced with si.*Control* or si.*PDGFRB* with mouse brain capillaries (Magenta) after 24 h incubation with brain slices (**c**), and quantification of iPSC-PC that homed to capillaries and extended processes along vessels, homed without processes visible, and did not home to mouse brain capillaries from 3 independent experiments per treatment (**d**). Scale bar: 50 µm. (**e**) Schematic showing proximity-dependent Proximity ligation Assay (PLA) for protein interactions between iPSC-PC PDGFRB and endothelial (EC) PDGF-BB, and between iPSC-PC vitronectin (VTN) and EC integrin α5 (ITGA5). (**f**) High-magnification of representative confocal imaging of proximity interactions (red) between PDGFRB and PDGF-BB (left) and between VTN and ITGA5 (right) in CellTracker labeled iPSC-PC (green) and lectin-positive brain capillaries (magenta) after 24 h incubation with brain slices. Merged images show orthogonal views. Data in **d** are stack bars with mean ± SEM; significance by Šídák’s multiple comparisons test between each homing category.

Then we asked whether iPSC-PC can home to brain capillaries in host brains with reduced pericyte coverage. To address this question we studied integration of CellTracker-labeled iPSC-PC with brain capillaries of 6-8-month old pericyte-deficient *Pdgfrb^F^*^7^*^/F^*^7^ (F7/F7) mice that exhibit approximately 40% loss of pericyte coverage ^3,23^. First, we used an *in vitro* live brain slice model (**Fig. 4a**) in which iPSC-PC were seeded onto the top of brain slices and allowed to integrate with the tissue over 24 h. Since pericyte-endothelial PDGFRB/PDGFB signaling is a key mechanism by which pericytes are recruited to the vessel wall ^24^, we also studied whether silencing PDGFRB in iPSC-PC compared to si.*Control* (**Fig. 4b**) will have an effect on their integration with F7/F7 brain capillaries. This experiment revealed that within 24 h after adding iPSC-PC to brain slices 70% of iPSC-PC transduced with si.*Control* homed to the host brain capillaries to form hybrid vessels with elongated processes similar to native, endogenous pericytes, whereas 20% associated with vessels without processes (**Fig. 4c**, **d**). After PDGFRB silencing, only 18% iPSC-PC homed to capillaries and exhibited elongated processes (p < 0.001) (**Fig. 4c**, **d**), whereas approximately 50% failed to home when compared to only 10% in control iPSC-PC (p < 0.001) (**Fig. 4c**, **d**).

To visualize pericyte-vasculature interaction in more detail, we performed high-resolution 3D rendering of representative iPSC-PC, showing enhanced elongation and homing to capillaries in si.*Control* transduced compared to si.PDGFRB transduced-iPSC-PC (**Supp. Video 1-2**). Additionally, we confirmed a functional association between iPSC-PC and the host vasculature using a proximity ligation assay (PLA), which detects protein colocalization within 40 nm proximity, with two previously described pericyte-endothelium protein-protein/ligand-receptor molecule pairs: PDGF-BB/PDGFRB^25^ and vitronectin/ Integrin α5 (ITGA5)^5^ (**Fig. 4e**). The expression of both functional protein pairs was detected specifically at sites of overlap between CellTracker labeled iPSC-PC and Lectin-stained vessels, indicating their close functional association within the formed hybrid brain vessels (**Fig. 4f**).Importantly, non–vessel-associated iPSC-PCs did not show any detectable PDGF-BB/PDGFRB or vitronectin/Integrin α5 PLA signal, confirming that proximity labeling occurred exclusively at vessel contact sites (**Supp. Fig. 5**).

**Figure 5.**
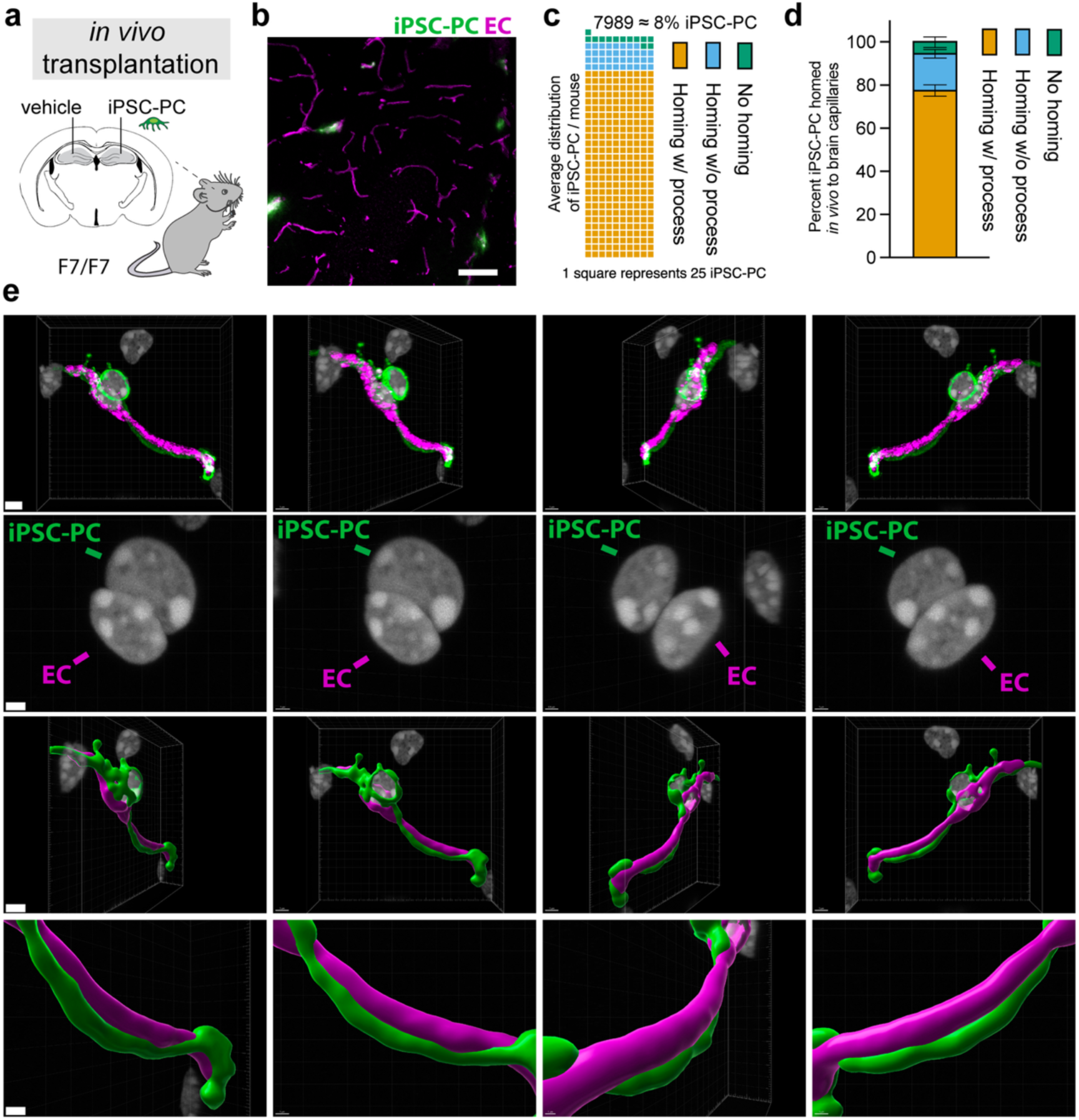
Transplanted iPSC-PCs associate with mouse microvessels. (**a**) Schematic of the in vivo experimental setup. Vehicle (aCSF) or 10⁵ CellTracker-labeled iPSC-derived pericytes (iPSC-PCs) were stereotaxically injected into the left (contralateral) and right (ipsilateral) hippocampus of 6–8-month-old Pdgfrb^F7/F7^ (F7/F7) mice, respectively. (**b**) Representative confocal image showing iPSC-PCs (green) aligned with endothelial cells (ECs, magenta). (**c**) Quantification of surviving iPSC-PCs classified as homing with processes, homing without processes, or not homing to vessels. Each square represents 25 cells. (**d**) Proportional distribution of iPSC-PCs across the three categories 48 h post-transplantation. (**e**) High-resolution 3D renderings of representative iPSC-PCs (green) wrapping around ECs (magenta). Sequential rotations show fluorescence signals and surface reconstructions at high magnification, highlighting iPSC-PC nuclei (gray) and elongated vessel-associated processes. Scale bars: 3 µm (overview: row 1,3), 1 µm (zoom-ins, row 2,4). n= 8 mice in **c,d.** Data in **d** are stack bars with mean ± SEM.

To assess if the homing effect can be also seen *in vivo,* we next transplanted iPSC-PC into the hippocampi of F7/F7 mice (**Fig. 5a**). The majority of transplanted iPSC-PCs successfully homed to brain capillaries and extended elongated processes along the vessel wall, as shown by high-resolution z-projections (**Fig. 5b-e, Supp. Fig 6**) and sequential sections confirming close spatial association with endothelial cells (**Supp. Fig. 7**). Consistent with the ex vivo results, approximately 75-90% of transplanted iPSC-PCs observed associated with vessels and displayed an elongated morphology, whereas only a small fraction remained non-vessel associated (**Fig. 5c, Supp. Fig. 8**). Based on quantitative analysis across serial sections, we estimate that ∼8% of all transplanted iPSC-PCs successfully engrafted and integrated with host brain capillaries (**Fig. 5d, Supp. Fig. 8**), which is within the expected range of similar transplantations^26,27^. High resolution 3D renderings further confirm close contact of iPSC-PCs (green) with endothelial cells (magenta) with intact nuclei, forming continuous hybrid vessel structures (**Fig. 5e, Supp. Video 3-5**).

**Figure 6.**
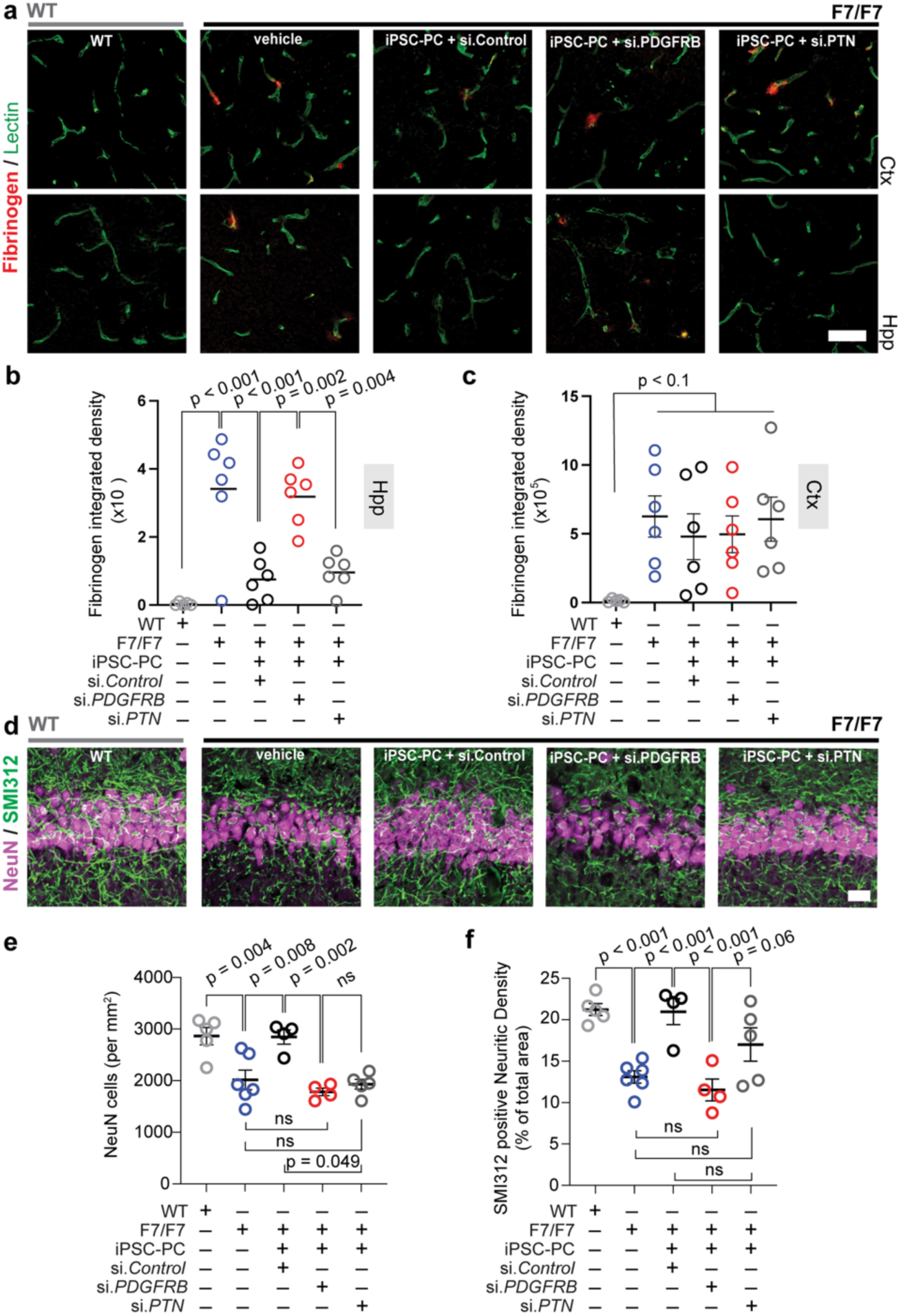
iPSC-PC transplantation reduces BBB leakage and improves neuronal retention in the hippocampus of pericyte-deficient mice. (**a-c**) Representative confocal microscopy imaging of extravascular fibrinogen and lectin-positive vasculature in the hippocampi and cortices in F7/F7 mice 5 days after vehicle or transplantation of iPSC-PC transduced with *si.Control*, *si.PDGFRB* or si.*PTN*, and wild type (WT) control mice (control) (**a**), and quantification of integrated extravascular fibrinogen signal in the hippocampus (**b**) and the cortex (**c**). Scale bar in (**a**) is 25 µm (**d-f**) Representative confocal microscopy images (**d**), and quantification of NeuN+ neurons (magenta; **e**) and SMI312+ neurofilaments (green; **f**) in the CA1 hippocampus in control mice and F7/F7 mice 5 days after vehicle or transplantation of iPSC-PC transduced with *si.Control*, *si.PDGFRB* or si.*PTN.* Scale bar in (**d**) is 50 µm. Data in **b**, **c**, **e**, **f** are mean ± SEM; n= 4 control mice in **b**,**c**, and n=5-6 mice for all other conditions in **b**, **c**, **e**, and **f**. Circles represent individual mice. Significance by one-way ANOVA followed by Tukey’s or Dunnett’s posthoc test.

**Figure 7:**
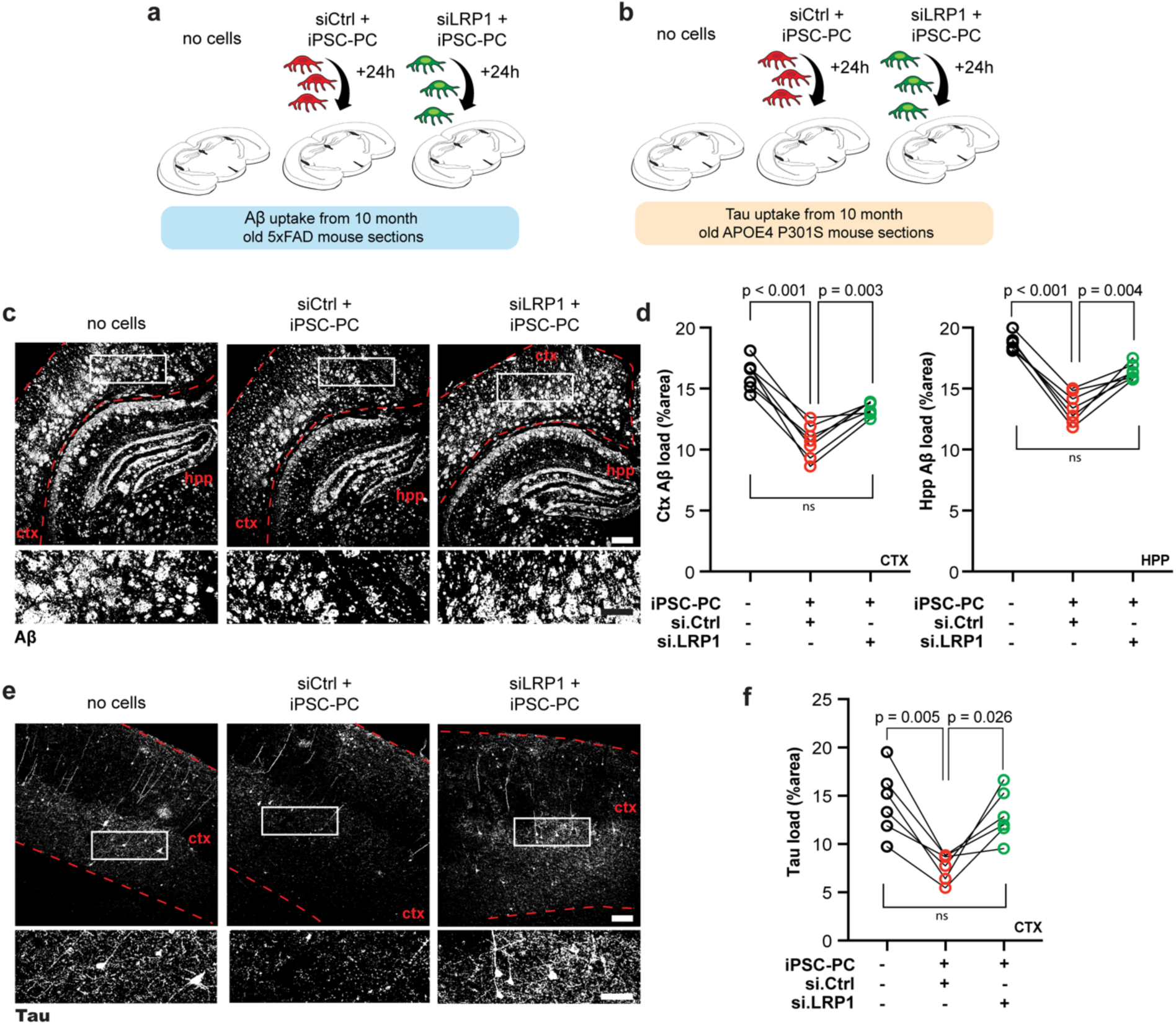
Aβ and tau clearance by iPSC-PC in 5xFAD and APOE4 P301S mouse brain sections is LRP1 dependent. **(a-b)** Schematic representation of siCtrl + iPSC-PC, siLRP1 + iPSC-PC, and no-cell control groups incubated with brain sections from 10-month-old 5xFAD mice (**a**) and 10-month-old APOE4 P301S mice (**b**). (**c-d**) Representative confocal images showing amyloid (white) in 5xFAD tissue sections after 24 h incubation with iPSC-PC + *siCtrl*, iPSC-PC + *siLRP1*, or control sections incubated with media alone (**c**), and quantification of amyloid load in the cortex and hippocampus in all groups (**d**). In (**c**), boxes show locations of insets. (**e-f**) Representative confocal images showing showing tau (white) in APOE4 P301S tissue sections after 24 h incubation with iPSC-PC + *siCtrl*, iPSC-PC + *siLRP1*, or control sections incubated with media alone (**e**), and quantification of cortical tau load in all groups (**f**). In (**e**), boxes show locations of insets. Data in **d**, **f** represent paired consecutive sections from the same mouse brain, with pairs connected by lines; n = 6-7 sections. Each circle represents an individual section. Significance was assessed by a repeated measure one-way ANOVA with Tukey’s multiple comparison test. Scale bars: 200 µm (**c,e** overview), 50 µm (**c,e**, insets).

We next sought to determine if transplanted iPSC-PC can restore functionally the BBB integrity in hippocampi of 6-8-month old F7/F7 mice, which exhibit substantial BBB leakage consequent to pericyte loss ^3,4^. To assess vascular integrity, we evaluated extravascular fibrinogen, a blood-derived protein that normally does not cross the intact BBB, in the hippocampus (the iPSC-PC graft site) and the cortex (a region distant from the site of iPSC-PC injections). This experiment confirmed that control F7/F7 mice treated with vehicle have higher fibrinogen extravasation immunofluorescence compared to age-matched controls on the same genetic background (p < 0.001), indicating a leakier BBB as previously reported ^3,4^ (**Fig. 6a-b**). However, F7/F7 mice transplanted with iPSC-PC transduced with si.*Control* showed substantially improved BBB permeability in the hippocampus, similar to values found in normal control mice on the same genetic background (**Fig. 6a-b**). The restoration of BBB integrity was seen in the hippocampus, a site of iPSC-PC grafting, but did not extend to regions distant from the grafting site such as cortex (**Fig. 6 a,c**). We then confirmed that restoring BBB permeability was dependent on the functional interaction PDGFB-PDGFRB. We determined that silencing of PDGFRB abolished the protective effects of iPSC-PC on BBB integrity, as fibrinogen leakage remained elevated, similar to vehicle treated F7/F7 mice (**Fig. 6 a-b**).

To determine whether grafting of iPSC-PC can influence neurodegenerative process in F7/F7 mice we stained hippocampal tissue with NeuN and SMI312 and determined neuronal counts and neuritic density, respectively, as previously reported ^4,28^ (**Fig. 6d-f**). Consistent with previous reports, 6-8-month old F7/F7 mice displayed 27% lower neuronal counts and 40% lower neuritic density compared to age-matched wildtype control mice on the same genetic background ^3^ (**Fig. 6d-f**). Treatment of F7/F7 mice with iPSC-PC transduced with si.*Control* had a substantial neuroprotective effect, as shown by 41% increase in NeuN-positive cells and 65% increase in SMI312 neuritic density in the CA1 region of the hippocampus, which was abolished by PDGFRB silencing (**Fig. 6d-f**). This data suggests that iPSC-PC can not only restore BBB integrity but also provides neuroprotection.

Since pleiotrophin (PTN), a neurotrophic factor secreted by brain pericytes, supports neuronal survival ^4,29^, we next asked whether silencing PTN has an effect on neuroprotective activity of iPSC-PC. First, we found that PTN silencing (**Supp. Fig. 9a**) did not result in loss of vasculoprotective effect of iPSC-PC in F7/F7 mice (**Fig. 6a-c**) consistent with a previous report showing that intracerebroventricular infusion of PTN does not affect vasculature in mice with inducible pericyte loss^4^. However, grafting of iPSC-PC transduced with si.*PTN* compared to si.*Control* in F7/F7 mice diminished the number of NeuN-positive neurons by 33% (p < 0.05) but not the SMI312-positive neuritic density (18.3% reduction, p = 0.32; **Fig. 6d-f**). These findings suggest that iPSC-PC can promote neuronal survival not only by restoring the BBB integrity but also partially through PTN-mediated neuroprotection, although PTN-independent mechanisms may also contribute to restore neuritic density.

Since mouse pericytes have been shown to clear Aβ ^7^ and tau ^8^ from brain, we also studied whether our iPSC-PC are capable of clearing Aβ and tau, and whether clearance is dependent on lipoprotein receptor LRP1, a major Aβ clearance receptor ^30,31^. First, iPSC-PC with and without LRP1 silencing (**Supp. Fig. 10a**) were seeded onto coverslips coated with Cy3-labeled Aβ42 (**Supp. Fig. 10b**), as we reported ^32,33^. After 5 days of incubation with iPSC-PC, we found that approximately 67% of coverslip-bound Cy3-Aβ42 was cleared compared to coverslips without cells (**Supp. Fig. 10c,d**). Aβ42 clearance was reduced by 64% after *LRP1* silencing (**Supp. Fig. 10c,d**), but was not affected by silencing other lipoprotein receptors *LDLR*, *VLDLR*, and *APOER2* (**Supp. Fig. 9b, 10d**). iPSC-PC were also seeded onto blank coverslips and incubated with soluble Cy3-labeled tau. After 48 hours, Cy3-tau was internalized and degraded by iPSC-PC (**Supp. Fig. 10e,f**). Silencing LRP1, which acts as a master regulator of tau uptake ^34^, significantly reduced Cy3-tau uptake in iPSC-PC by 74% (**Supp. Fig. 10e,f**). Importantly, we additionally demonstrate that iPSC-PCs begin to internalize Aβ and tau already within 24 hours, as shown by colocalization of fluorescently labeled Aβ42 and tau with LysoView-positive lysosomes, indicating rapid endocytic uptake and degradation (**Supp. Fig. 11**).

To further evaluate the ability of iPSC-PC to clear Aβ and tau in a pathological AD brain environment, we incubated CellTracker-labeled iPSC-PC for 24h on brain sections from 10-month-old 5xFAD mice, which develop a strong amyloid pathology^35^ (**Supp. Fig 12a-c**, **Fig. 7**), and APOE4-P301S mice, which carry the *APOE4* AD-risk allele and the P301S tau mutation to model strong tau aggregation^36^ (**Supp. Fig. 12 d-f, Fig. 7**). iPSC-PC efficiently internalized Aβ aggregates in cortical and hippocampal regions of 5xFAD brain sections (**Supp. Fig. 12b**), leading to a significant reduction in amyloid load compared to consecutive control sections incubated in media without iPSC-PC (p < 0.01, **Supp. Fig. 12c**). Similarly, when iPSC-PC were incubated on APOE4-P301S brain sections, we observed robust uptake of tau aggregates (**Supp. Fig. 12d-f**) compared to control sections (p < 0.01, **Supp. Fig. 12f**). To determine whether this clearance is LRP1-dependent, iPSC-PC were transduced with siLRP1 or siControl and incubated on consecutive brain sections (**Fig. 7a,b**). While si.Control iPSC-PC efficiently reduced Aβ (**Fig. 7c-d**) and tau levels (**Fig. 7 e-f**), LRP1 knockdown abolished these beneficial effects (**Fig. 7c-f**). These findings suggest that iPSC-PC effectively clear Aβ and tau in an *ex vivo* neurodegenerative AD environment and that this process is dependent on the integrity of the LRP1-dependent pathway, similar to previously described mouse pericytes ^7,8^.

## Discussion

Earlier characterization of iPSC-PC and primary brain PC relied on a relatively limited set of 4-10 pericyte markers and/or proteins known to be expressed by pericytes ^11–13^. While one study suggested parallels between iPSC-PC and brain PC using bulk RNAseq, the work focused on transcript presence or absence, leaving the statistical analysis of differential gene expression unexplored ^12^. Moreover, these previous studies ^11–13^ compared iPSC-PC to commercial human ‘primary brain pericytes’ of unspecified origin, likely derived from human fetal tissue. A fetal pericyte origin is also suggestive from the high expression of ACTA2, which has not been observed in iPSC-PC and PC in the present study, or in pericytes from two recent single-cell atlases of the human adult brain vasculature ^37,38^, but rather is found in immature pericytes ^39^. In contrast, here we performed the first comprehensive analysis of global protein expression and protein phosphorylation in iPSC-PC and PC, which revealed highly shared proteome with preserved functional characteristics of proteins as indicated by their comparable phosphorylation signatures, and with preserved multiple physiological functions of pericytes as shown by different experimental models.

Using a quantitative LC/MS comparative analysis of 8,344 proteins and 20,572 phosphopeptides, we showed that human iPSC-PC share 96% of total proteins and 98% of protein phosphopeptides with primary human brain PC including canonical pericyte markers, cell adhesion and tight junction proteins, transcription factors, and the protein kinase families of the human kinome. It has been reported that prolonged cultivation of PC (up to 20 passages) can alter gene expression, reducing extracellular matrix (ECM)-related genes ^40^. In our study, PC were sub-cultured for 2-4 passages before proteomic analysis due to the relative cell scarcity at collection from human brains. Our proteomic screen showed consistent ECM protein expression between iPSC-PC and PC, including integrins (Itga1, Itga4, Itga7, Itgb5), laminins (Lama2, Lama4, Lamb2, Lamc3) and collagens (Col4a1, Col4a2). While this suggests minimal culture-induced proteomic alterations, we cannot entirely exclude the possibility of subtle expression changes in PC due to sub-culturing. We show that iPSC-PC preserve BBB integrity through molecular and structural features of tight junctions, in agreement with prior studies reporting strengthened barrier function and increased TEER in endothelial co-cultures.^11,12^

We showed that iPSC-PC home to brain capillaries of pericyte-deficient F7/F7 mice on *ex vivo* brain slices, and *in vivo* after grafting into the hippocampus suggesting their vasculogenic potential to form hybrid vessels with the host vessels, which we show requires intact PDGFRB signaling mechanisms. Importantly, grafting of iPSC-PC into the hippocampus restores BBB permeability to control levels in F7/F7 mice. Moreover, transplantation of iPSC-PC into the F7/F7 mouse hippocampus increased neuronal counts and neuritic density. This neuroprotective effect of iPSC-PC required PDGFRB and was partially dependent on the neurotrophic factor PTN, suggesting other pericyte-mediated neuroprotective pathways may also be involved including e.g. other neurotrophic factors ^29,41^. Finally, we showed that iPSC-PC clear Alzheimer’s Aβ and tau proteins via LRP1 *in vitro* and on brain sections of aged 5xFAD and APOE4-P301S mice, similar as previously shown in mice ^30,31,34^. While our study demonstrates clearance of Aβ and tau aggregates in *ex vivo* experiments, future studies will need to assess whether iPSC-PC can exert similar Aβ and tau clearance effects longitudinally *in vivo* in AD models, particularly in relation to long-term graft survival and functional impact on AD pathology. Moreover, determining whether iPSC-PCs primarily mediate Aβ and tau clearance at the vascular interface or also within the parenchyma will be important for defining the underlying mechanisms of action *in vivo*.

In conclusion, the present study provides support for potentially translating iPSC-PC for use in humans to restore impaired cerebrovascular and brain functions. In this regard, the autologous patient-derived iPSC-PC brain transplants may hold potential as a valuable cell replacement therapy for neurological disorders associated with pericyte deficiency such as for example AD, vascular cognitive impairment, and related disorders ^6^ characterized by cerebrovascular changes and BBB breakdown, loss of neurons and impaired clearance of Alzheimer’s Aβ and tau neurotoxins.

## Materials and Methods

### Cells

#### Human iPSC

iPSC cultures were generated from skin fibroblasts from *e3/e3* donors, as previously reported ^42^ The cells were preserved in the liquid nitrogen vapor phase at −196 °C until differentiation.

#### Primary human PC

Primary human brain microvascular pericytes (PC) from **ε*3/*ε*3* donors were isolated from postmortem cortical tissue, as we previously described ^43^. First, brain capillaries were isolated, digested with collagenase, and the capillary fragments were seeded onto fibronectin-coated plates for 5-7 days. Next, cells were treated with 0.05% trypsin EDTA for 5 min to separate the detached PC from endothelial cells. PC were cultured in Earl’s modified essential medium containing 10% human serum, 20% newborn calf serum, 10 mM N-2-hydroxyethylpiperazine-N’-2-ethanesulfonic acid, 150 µg/ml endothelial cell growth factor, 5 U/ml heparin, 2 mM/L glutamine, and antibiotics for 2-4 passages, as previously described ^43,44^. PC expressed canonical pericyte markers PDGFRβ, NG2, and CD13 and did not express endothelial markers (CD31, GLUT1), astrocyte markers (GFAP, S100B, Aldh1l1), microglial markers (CD11b, IBA1, CD68), neuronal markers (NF-L, TUJ1), and oligodendrocyte markers (CNPase, Olig2) or epithelial markers, as we previously reported ^43^ and shown in **Supp. Fig. 2**. PC remained frozen in the liquid nitrogen vapor phase until they were cultured. Early passage (P2-P4) PC were cultured in a human pericyte medium (ScienCell, 1201) and used for liquid chromatography-mass spectrometry (LC-MS) analysis and Western blotting.

#### Other primary cells and cell lines (BEC, VSMC, SH-SY5Y, CRL-2266)

Primary human brain microvascular endothelial cells (BEC) were obtained and isolated from autopsies of neurologically normal individuals, as we described before ^32,45,46^. BEC (>98%) expressed endothelial markers vWF and CD105 and were negative for the microglial marker (CD11b), astrocyte marker (GFAP), and vascular smooth muscle cell marker (α-smooth muscle actin). Early passage cultures (P2-4) were used. BEC were frozen in the liquid nitrogen vapor phase (−196 °C) until they were cultured in a human endothelial cell medium (ScienCell, 1001). Human cerebrovascular smooth muscle cells (VSMC) were obtained and isolated from pial cerebral arteries (>100 μm diameter) and characterized as previously reported ^46,47^. VSMC (>98%) expressed α-actin, myosin heavy chain, calponin, and SM22α and were negative for endothelial cell markers (vWF and CD31), astrocyte marker (GFAP), and microglial marker (CD11b). Early passage (P2-4) cultures were used in the study. VSMC were cultured in human smooth muscle cell medium (ScienCell, 1101). Immortalized human neuronal cell line (SH-SY5Y, CRL-2266), human microglial cell line (HMC3, CRL-3304), and human oligodendroglioma cell line (BT88, CRL-3417) were purchased from ATCC and cultured according to ATCC instructions. Human astrocytes were obtained from ScienCell (CP1800). Cell pellets of all primary cells and cell lines were prepared for Western blotting.

#### In vitro BBB assay

Primary mouse brain endothelial cells (mBEC) were purchased from Cell Biologics, C57-6023, primary human brain endothelial cells (hBEC) were used from brain autopsies of young neurologically normal individuals, as we previously described ^43^. Cells were cultured in endothelial cell medium (Cell Biologics, M1168) on fibronectin-coated (Neuvitro, GG-12-1.5-Fibronectin) or gelatin-coated coverslips. For gelatin coating, coverslips (Fisher, 08-774-383) were incubated with 0.1% gelatin (Sigma, G1393) for at least 30 minutes at 37°C.

To investigate tight junction (TJ) proteins ZO-1 and claudin-5, hBEC and mBEC monolayers were treated with either iPSC-pericyte (iPSC-PC)-derived conditioned media or control endothelial cell medium. Conditioned medium was generated by culturing iPSC-PC in human pericyte medium (ScienCell, 1201) for 24 hours, followed by centrifugation at 500g to remove cells and debris.

For claudin-5 analysis, conditioned or control medium was applied to mBEC and hBEC cultures for 24 hours. For ZO-1 analysis, the conditioned medium was diluted 1:1 with fresh endothelial medium and applied for 48 hours. Control cultures received fresh endothelial medium only. Following treatment, monolayers were fixed with 4% paraformaldehyde for 10 minutes and stained for TJ markers (see Supp. Table 5 for antibodies), with nuclear counterstaining using either DAPI or DRAQ5 (1:1000, Novus Biologicals, NBP2-81125).

Images were acquired using a Nikon A1R confocal microscope with a 20x objective and NIS-Elements software. For quantification of TJ length, images were preprocessed using the median filter in Fiji, binarized with adaptive thresholding, and skeletonized to measure total TJ length. TJ length was normalized to the number of DAPI- or DRAQ5-positive nuclei. Calculations were automated, eliminating human bias.

### iPSC-PC differentiation via NCC intermediates

iPSC-PC were generated via NCC intermediates following previously reported protocols ^11,12^ with modifications. Briefly, iPSC were maintained on growth factor reduced matrigel (Corning, 356230) in mTeSR Plus (StemCell Technologies, 100-0276), and NCC were derived from iPSC using STEMdiff^TM^ Neural Crest Differentiation Kit (StemCell Technologies, 08610). Next, iPSC were washed with PBS, dissociated with Accutase (Thermo Fisher, A1110501) for 5 min, centrifuged at 200xg for 4 min, counted using a Countess 3 (Thermo Fisher, AMQAX2000) and seeded at a density of 0.75-1×10^5^ cells/cm^2^ in mTeSR plus (StemCell Technologies, 1000276) supplemented with 10 μM Y-27632 (Tocris, 1254). For each iPSC line seeding densities were adjusted to reach 100% confluency at differentiation day 3-4. Media changes were performed daily using NCC media from the STEMdiff kit. After 6 days of differentiation, iPSC-derived NCC were dissociated using Accutase incubation for 5 min and purified using EasySep^TM^ release human PSC-derived NCC positive selection kit (StemCell Technologies, 100-0047) as per manufacturer’s instructions. Following purification, cells were replated onto Matrigel-coated surfaces in NCC media supplemented with 10 μM Y-27632. The purified NCC were replated onto glass coverslips for imaging, and on day 7, NCC coverslips were fixed and stained according to the immunocytochemistry methods described below. Imaging of the NCC revealed expression of typical neural crest markers SOX10 and p75^NTR^.

To further differentiate to iPSC-PC, iPSC-derived NCC were replated onto matrigel-coated 6-well plates at 1:2 split ratio, and pericyte medium (ScienCell, 1201) was added the next day. Media were changed daily, and once cells reached 70-80% confluence, they were passaged using 0.05% Trypsin/EDTA (Thermo Fisher, 25300054) and replated onto poly-L-lysine (PLL)-coated dishes. After 14 days, differentiated iPSC-PC were cultured onto glass coverslips for imaging. iPSC-PC were positive for the canonical pericyte markers PDGFRβ, CD13, NG2, Caldesmon, and Vitronectin, and were negative for the BEC markers (CD31, GLUT1), astrocyte markers (GFAP, S100B, Aldh1l1), microglial markers (CD11b, IBA1, CD68), neuronal markers (NF-L, TUJ1), oligodendrocyte markers (CNPase, Olig2), and VSMC markers (myosin heavy chain/MYH11, α-smooth muscle actin/ACTA2, calponin/CNN1), and epithelial markers (CDH1, KRT18, EpCAM). iPSC-PC were maintained on PLL-coated dishes, and media were changed every other day.

#### Immunocytochemistry for cell culture

iPSC-derived NCC, iPSC-PC, PC, and mouse BEC (Cell Biologics, C57-6023) were plated and cultured on glass coverslips. Cells were rinsed twice with sterile PBS and then fixed with 4% PFA for 10min. Next, 3x 5 min washes in PBS were performed before blocking in PBS containing 5% donkey serum and 0.5% Triton X-100, and blocked for at least 1 h. NCC, iPDC-PC, and PC were incubated overnight with SOX10, p75, PDGFRβ, and/or NG2 primary antibodies (listed in Supp. Table 5) diluted in PBS containing 5% donkey serum (PBSD). On the next day, 5x 8-min washes in PBS were performed and cells were incubated with secondary antibodies (listed in **Supp. Table 5**) in PBSD for 1-2 h. After 4x 8-min washes in PBS, coverslips were mounted in a DAPI-containing mounting medium (Southern Biotech, 0100-20) and imaged on a Nikon A1R confocal microscopy system with NIS-Elements software control using a 20x objective.

#### Quantification of PDGFRB- and NG2-positive cells

All samples were stained and imaged using identical image acquisition settings; images were processed and analyzed using ImageJ (Fiji), similar to previous studies ^23^. Fluorescence signal from the marker of interest (PDGFRB, NG2) was thresholded using Otsu thresholding plugin ^48^ in each image. The thresholding was performed in the same manner for all samples. Analyze Particles function was used to determine the number of marker-positive cells. To calculate the percentage of positive cells, the number of marker-positive cells was normalized to the total nuclei count for each image.

### Genetic manipulation of iPSC-PC

#### Generation of GFP-expressing iPSC-PC

GFP-iPSC-PC were generated by dissociating the cells with 0.05% trypsin-EDTA, resuspending in 200 µL media, and then incubating with either lenti-PGK-GFP or lenti-CMV-GFP (SignaGen SL100268) at a multiplicity of infection (MOI) ranging from 1:1 to 10:1 (according to the manufacturers recommendation) for 45 min at 37°C along with 10 µg/mL polybrene (Sigma, TR1003G) to enhance transduction efficiency. To produce lenti-PGK-GFP, 293T cells were co-transfected with a lentiviral vector (Open Biosystems, RHS4696-99634943) together with VSVG expression vector (Addgene 8454) and gagpol expression vector (Addgene 14887). After 12-16 h, the media was discarded and replaced with serum-free Ultraculture media (Lonza, 12-725F) with 2X glutamine for virus harvesting. Viral supernatant was collected the next day and concentrated by spinning at 15,000 rpm at 4°C overnight in an ultracentrifuge. The virus was resuspended in the remaining media by shaking on ice for 2-12 h and added to 200 µL cell suspension at a volume of 50-150 µL. PGK-GFP expressing iPSC-PC were used to study grafting into the hippocampus *in vivo* which included fibrinogen extravasation measurements of BBB permeability and tissue analysis of neuronal changes. PGK-GFP and CMV-GFP expressing iPSC-PC were used in clearance assays to evaluate the ability of iPSC-PC to clear Alzheimer’s Aβ and tau neurotoxins.

#### PDGFRB and PTN silencing in iPSC-PC

To silence PDGFRB, 70-80% confluent iPSC-PC were treated with either Dharmacon AccellTM SMARTPool human PDGFRB siRNA (si.*PDGFRB*) (Dharmacon, E-003163-00-0010), Dharmacon AccellTM SMARTPool human PTN siRNA (si.*PTN*) (Dharmacon, E-018793-00-0020) or AccellTM non-targeting control pool siRNA (si.*Control*) (Dharmacon, D-001910-10-50) at a concentration of 1 µM in Dharmacon Accell siRNA delivery media (Dharmacon, B-005000-500) in 6-well culture plates for 48 h per the manufacturer’s instructions. The efficacy of PDGFRB and PTN silencing was confirmed by Western blotting. si.*Control* and si.*PDGFRB* iPSC-PC were used in studies on brain slices and for transplantation into the hippocampus of pericyte-deficient mice (see below). si.*PTN* iPSC-PC were used for *in vivo* transplantation in pericyte-deficient mice (see below).

### Pericyte Contractility Assay

iPSC-PC and PC contraction was assessed using KCl stimulation. Glass-bottom dishes (Cellvis, D35C4) were coated with poly-D-lysine (PDL; Sigma) and seeded with 10,000 iPSC-PC or PC per compartment. On the day of the assay, culture medium was replaced with 200 µL pre-warmed HBSS (Thermo Fisher), followed by 200 µL HBSS with 150 mM KCl (final 75 mM). Time-lapse imaging (3 frames/min) was performed for 30 min using an inverted confocal fluorescence microscope (Nikon A1R confocal with NIS-Elements software). Pericyte length was measured along the longest axis at 0 and 20 min using ImageJ, and contraction was calculated as the percent change in cell length.

### Western blotting

The cell pellets of primary human iPSC-PC, PC, BEC, VSMC, primary astrocytes (ScienCell, CP1850), and immortalized human neuronal cell line (SH-SY5Y, ATCC, CRL-2266), microglial cell line (HMC3, ATCC, CRL-3304), oligodendroglioma cell line (BT88, ATCC, CRL-3417) and human epithelial cells (Cell Biologics, H-6034) were resuspended in RIPA buffer (50 mM Tris pH 8.0, 150 mM NaCl, 1% NP40, 0.1% SDS, 0.5% sodium deoxycholate and Roche protease inhibitor cocktail) and sonicated. Then, samples were centrifuged for 30 min at 20,000 x g, and supernatants used for protein quantification (Thermo Fisher, 23,228). Samples were prepared with NuPAGE LDS sample buffer (Invitrogen, NP0007) and a sample reducing agent (Invitrogen, NP0009), heated at 70 °C for 10 min. Proteins (5–10 μg protein loaded per sample) were separated by electrophoresis on NuPAGE Novex Bis-Tris precast 4–12% gradient gels (Thermo Fisher, WG1402BOX). Following electrophoretic transfer, nitrocellulose membranes were blocked using a blocking buffer (Thermo Fisher, 37,536) and incubated overnight at 4 °C with primary antibodies diluted in a blocking solution. After washing with tris buffered saline containing 0.1% Tween 20 (TBST), membranes were incubated with horseradish peroxidase (HRP)-conjugated donkey anti-rabbit, anti-mouse, or anti-goat secondary antibody for 1 h at room temperature (see Supp. Table 6 for details), rewashed in TBST and treated for 5 min with Super Signal West Pico chemiluminescent substrate (Thermo Fisher, 34580) or Pierce ECL Western blotting chemiluminescent substrate (Thermo Fisher, 32106). Membranes were exposed to CL-XPosure film (Thermo Fisher, 34091) and developed in MINI-MED 90 X-ray film processer (AFP Manufacturing corporation, Peachtree City, GA).

### Proteomics analysis

A quantitative comparative analysis of total proteome and phosphoproteome of six separate cultures from three iPSC lines (iPSC-PC 1-6) and six cultures of PC from three patients (PC1-6) was performed using LC-MS, as we previously described ^9,16–19^.

#### Sample preparation

400,000 cells per sample were dissolved in 100 µL lysis buffer (0.5 M triethylammonium bicarbonate, 0.05% sodium deoxycholate) and sonicated (Q700, QSonica, amplitude = 10, 2 sec on/2 sec off pulses, 20 sec total processing time per sample, on ice) and then centrifuged at 15,000 rpm at 4°C for 10 min. The supernatant was transferred on ice to a fresh tube and the protein concentration determined using the Qubit Protein Assay Kit (Thermo, Q33211) and Qubit 4.0 fluorometer per manufacturer’s instructions. Equal amount of protein (40 µg) per sample was transferred to a fresh tube adjusted to highest volume (90 µL) with lysis buffer. Next, 4 µL Reducing Reagent (Sigma, 4381664) were added to each sample and samples were incubated at 60°C for 1hr. Next, 2 µL Alkylating Reagent (Sigma, 4381664) were added to each sample and samples were then incubated at room temperature for 15 min. Lastly, 2 µg trypsin/LysC (Promega, V50703) were added to each sample. Samples were incubated overnight at room temperature in the dark. TMTpro reagents (Thermo, A34808) were equilibrated and 20 µL of anhydrous acetonitrile (Sigma, 900644) were added to each label. The solution was then transferred to each sample and incubated at room temperature for 1 h. Next, 8 µL 5% hydroxylamine were added to each sample. Samples were incubated for 15 min. Samples were then combined per multiplex experiment and dried-up using a speedvac (Eppendorf 5301 vacufuge concentrator). The high-select TiO2 phosphopeptide enrichment kit (Thermo, A32993) and high-select Fe-NTA phosphopeptides enrichment kit (Thermo, A32992) were sequentially used per manufacturer’s recommendations. All steps were performed at room temperature.

#### Quantitative proteomic analysis with LC-MS

The labelled native peptides fractionated using offline alkaline reverse phase nano-chromatography. Fractions were dried-up up using a speedvac (Eppendorf 5301 vacufuge concentrator), reconstituted in ddH2O 0.1% formic acid and analyzed using LC-MS with FAIMS ion mobility pre-separation (nano-easy LC 1200, Thermo Orbitrap Exploris 480).

#### Data processing

Raw data files of the native peptide LC-MS analysis were submitted to Proteome Discoverer 2.5 (Thermo) for target decoy search using Sequest against the homo sapiens canonical swissprot database (TaxID= 9606, v2021-10-30). The search allowed for up to two missed cleavages, a precursor mass tolerance of 20ppm, a minimum peptide length of six and a maximum of three equal dynamic modifications of oxidation (M), deamidation (N, Q) or phosphorylation (S, T, Y). Methylthio (C) and TMTpro (K, peptide N-terminus) were set as static modifications. Peptide level confidence was set at q<0.05 (<5% false discovery rate, FDR). Only unique peptides were considered for protein quantitation. Percent co-isolation excluding peptides from quantitation was set at 50. All proteins expressed in iPSC-PC and PC were normalized against the total amount of protein loaded in each sample. Protein level abundances were median normalized and scaled to derive their relative abundance similar to reported previously ^9^. Raw data files of the phosphopeptide LC-MS analysis were submitted to Proteome Discoverer 2.5 (Thermo) for target decoy search using Byonic against the homo sapiens canonical swissprot database (TaxID= 9606, v2021-10-30). The search allowed for up to two missed cleavages, a precursor mass tolerance of 20ppm, a minimum peptide length of six and dynamic modifications of oxidation (M) (common 1), deamidation (N, Q) (common 1), phosphorylation (S, T, Y) (common 2), TMTpro (K, peptide N-terminus) (common 2). Methylthio (C) was set as static modification. All phosphopeptides expressed in iPSC-PC and PC were normalized against the total amount of protein loaded in each sample. Protein level abundances were median normalized and scaled to derive their relative abundance similar to reported previously ^9^. All mass spectrometry proteomics data have been deposited to the ProteomeXchange Consortium via the PRIDE partner repository, accession: PXD043995.

#### Statistics for proteomics analysis

We calculated the mean relative abundance for all proteins and phosphopeptides in six iPSC-PC and six PC samples, as well as the ratios of the mean relative abundance for each protein and phosphopeptide in iPSC-PC vs. PC samples. Ratios were log2 transformed, and ratios > 1.35 and < 0.66 were considered significant as we previously reported ^16,17^. We performed a two-sample T-test (unpaired, heteroscedastic) to identify differentially expressed proteins between the two groups. The two-stage Benjamini Yekutieli Krieger step up method was used for multiple hypothesis testing FDR correction of the p-value. An FDR<0.01 (q < 0.01) was considered significant.

#### Data visualization

Parts of data visualization was additionally performed using R software (Verson 4.2). Heatmaps of commonly and differentially expressed protein z-scores were generated using pheatmap function. Other graphs including volcano plots, donut plots, and violin plots were generated using ggplot2 function in R, as previously reported ^49–51^.

### Mice

Platelet-derived growth factor receptor β mutant mice, *Pdgfrb^F^*^7^*^/F^*^7^, on 129S1/SvlmJ background were used for *ex vivo* and *in vivo* studies, as described below. These mice express PDGFRb with seven point mutations that disrupt specific signal transduction pathways; including residue 578 (Src), residue 715 (Grb2), residues 739 and 750 (PI3K), residue 770 (RasGAP), residue 1008 (SHP-2), by changing the tyrosine to phenylalanine, and residue 1020 (PLCγ), where tyrosine was mutated to isoleucine, as reported ^52^. *Pdgfrb^F^*^7^*^/F^*^7^ mice express mutant PDGFRβ exclusively in perivascular mural cells including pericytes, and not in neurons, astrocytes or brain endothelial cells ^3,53^. As control, we used age-matched 129S1/SvlmJ mice with two copies of the wild-type *Pdgfrb* gene. For the Aβ and tau clearance experiments, we used brain sections from 10-month old 5xFAD mice, which carry five familial Alzheimer’s disease mutations in APP and PSEN1, leading to strong amyloid-β deposition ^35^, and APOE4-P301S mice, which carry the human *APOE4* AD risk allele and the P301S tau mutation that exacerbates tau hyperphosphorylation and aggregation ^36^. The mice were housed in groups under standard conditions on a 12 h light cycle, and with ad libitum access to food and water. Both male and female mice 6-8-month old were used in the study. All procedures were approved by the Institutional Animal Care and Use Committee at the University of Southern California using US National Institutes of Health guidelines. All procedures followed the ARRIVE guidelines ^54^. The operators responsible for experimental procedure and data analysis were blinded and unaware of group allocation throughout all experiments.

### *Ex vivo* brain slice preparation and imaging analysis

*Ex vivo* brain slices were prepared similar to our previous studies ^55^ with modifications. Pdgfrb*^F^*^7/*F*7^ mice were retroorbitally injected with 80 μL DyLight-649 labeled Lycopersicon esculentum lectin (Vector Laboratories, DL-1178-1) ^3^ to label blood vessels at least 15 min prior to euthanasia. Mice were deeply anesthetized with 5% Isoflurane, then rapidly decapitated, and brains extracted into ice-cold Hibernate A minus Phenol Red medium (Transnetyx Tissue, HAPR500). Slice preparation was performed using a Leica VT1000S vibratome. Coronal slices (250 μm) containing the hippocampus were then cut in ice-cold Hibernate A minus Phenol Red medium.

After sectioning, slices were transferred to individual wells of a multiwell plate and maintained in a homing medium consisting of DMEM (Gibco, 10569010), 5% FBS (Gibco, 16140-071), Pen-Strep (Gibco, 15140122), 2% platelet lysate (Stem Cell Technology, 200-0323), an antioxidant (Sigma-Aldrich, A1345), GlutaMAX (Gibco, 35050061), Na-Pyruvate (Gibco, 11360070), CultureBoost (Cell Systems, 4CB-500), and Pericyte Supplement (ScienCell, 1252).

For pericyte homing assays, 50,000 iPSC-PC labeled with CellTracker were added on top of each brain slice and incubated for 24 h at 37°C with 5% CO₂. Brain slices were washed 3 times in PBS, fixed in 4% PFA for 30 min, mounted on slides, coverslipped with mounting media and imaged on a Nikon A1R confocal microscope system with NIS-Elements software control (20x objective, 1.1 numerical aperture). CellTracker-positive iPSC-PC were visualized throughout the brain slices and scored as (a) being associated with a vessel and forming processes, (b) associated with a vessel without processes or (c) not associated with a vessel. The percentage of homed iPSC-PC with/without processes to brain capillaries was calculated by dividing the number of homed iPSC-PC with/without processes with a vessel to the total number of iPSC-PC within all fields imaged in each brain slice.

### Proximity ligation assay (PLA)

For proximity ligation assay (PLA), NaveniFlex Tissue GR Red (Navinci, 39220) was used following the manufacturer’s protocol. Brain slices with homed iPSC-PC from the *ex vivo* brain slice preparation were fixed with 4% PFA for 30 min and then blocked and incubated overnight at 4°C with primary antibodies: rabbit anti-PDGF-B (Invitrogen, MA5-51346, 1:100) and goat anti-PDGFRβ (R&D Systems, BAF385, 1:50), or goat anti-vitronectin (R&D Systems, AF2349, 1:50) and rabbit anti-integrin α5 (Abcam, ab150361, 1:100) (**Supp. Table 5**). After washing in TBS-T (0.05% Tween-20), secondary PLA probes were applied, and signal amplification was performed per the kit instructions. PLA signals were imaged using confocal microscopy, and positive interactions were shown using z-projections in Fiji (ImageJ).

### *In vivo* transplantation of iPSC-PC and imaging analysis

Before transplantation, one well of GFP-iPSC-PC or CellTracker-labeled iPSC-PC were rinsed with PBS and incubated with 0.05% Trysin/EDTA (Thermo, 25300054) for 3 min to dissociate the cells. The cells were then centrifuged at 200xg for 4 min, and the resulting pellet was resuspended in 1 mL of pericyte medium for counting. The number of cells per well was determined using a Countess 3 (Thermo, AMQAX2000) per manufacturer’s instructions. A second well was then dissociated and centrifuged, and pellet was resuspended in appropriate volume of sterile artificial CSF solution (aCSF) (Harvard Apparatus, 597316) to achieve a concentration of 10^5^ cells/ mL. *Pdgfrβ^F^*^7^*^/F^*^7^ mice were anesthetized with 100 mg ketamine/10 mg xylazine per kg body weight, placed in a stereotactic apparatus (ASI Instruments) and fixed accordingly. During the procedure, rectal temperature was maintained between 36.5 and 37.0°C using a feedback-controlled heating system. A midline incision of approximately 1 cm was made through the skin to reveal the skull. A dental drill was then used under a surgery microscope to carefully drill a small hole on the bone over a desired brain nucleus. Vehicle (1 µL aCSF; left hippocampus) and iPSC-PC (10^5^ cells in 1 µL aCSF) transduced with si.*Control* or si.*PDGFRB* or si.*PTN* (right hippocampus) were stereotaxically injected into the dorsal hippocampus (coordinates A/P: −2.2; M/L: 1.7; and D/V: 1.8). All mice received daily immunosuppressive treatment with cyclosporine (Novartis, Sandimmune 10 mg/kg i.p.) for 24, 48 h or 5 days, as described in our previous study ^56^. We chose the hippocampus for iPSC-PC transplantation due to early pericyte degeneration and BBB leakage in AD, making it a relevant therapeutic target for pericyte-vascular interactions. Also, similar studies have also used pericyte transplantation into the hippocampus ^7^.

Mice were euthanized 24, 48 h and 5 days after iPSC-PC grafting. Animals were anesthetized with an i.p. injection of 100 mg/kg ketamine and 10 mg/kg xylazine and perfused transcardially with 30 ml of PBS containing EDTA. Brains were removed and embedded in O.C.T. compound (Tissue-Tek) on dry ice and cryo-sectioned at a thickness of 20 µm.

#### Quantification of iPSC-PC

For imaging iPSC-PC homing, mice received a 50 µl retroorbital injection of Lycopersicon esculentum DyLight 649-lectin (Vector Laboratories, DL-1178-1) 15 min prior to euthanasia to identify blood vessels. Sections were cover slipped using fluorescent mounting medium (Dako) and imaged with a Nikon A1R HD inverted confocal microscope with Galvano scanner using high resolution optical sections. At least 10 images containing iPSC-PCs were collected per mouse for quantification. Individual iPSC-PCs were classified based on their spatial relationship to lectin-positive vessels as either (1) vessel-associated with elongated processes, (2) vessel-associated without visible processes, or (3) non-associated. The proportion of cells in each category was calculated as a percentage of the total number of detected iPSC-PCs per mouse. To estimate the total number of iPSC-PC 48 h post-transplantation the brain, we first identified the section containing the highest density of grafted cells and used this to approximate peak cell graft abundance. A Gaussian distribution was then fitted across serial brain sections to estimate the spatial spread of the graft along the sectioning axis. To correct for potential double-counting of split nuclei, the Abercrombie correction was applied, and the resulting data were used to derive the total number of surviving cells per brain (approach similar to prior quantitative graft analyses.^57–59^

#### High-Resolution Confocal Microscopy

Imaging was performed using a Leica Stellaris 8 microscope (Leica Microsystems) equipped with a Leica scanhead, a white-light tunable laser (excitation range: 440–790 nm), and Leica HyD S and HyD X photon-counting detectors. A HC PL APO CS 93×/1.30 motCORR glycerol immersion objective was used. To visualize DAPI, a 405 nm laser was used for excitation, and emission was collected in the 430–480 nm range. For PC and EC visualization, the white-light laser was tuned to 488 nm and 638 nm, respectively, with a Leica ND filter applied. Emission was collected in the 500–560 nm range for PC and 650–790 nm for EC using the HyD S detector. The voxel size of the acquired images was 41 × 41 × 330 nm. Image processing and visualization were performed using Bitplane Imaris 10.2 (Oxford Instruments). Individual cell segmentation was carried out using Labkit for pixel classification.

### Immunohistochemistry

#### Tissue processing and imaging

The mice were anesthetized with an i.p. injection of ketamine (100 mg/kg) and xylazine (50 mg/kg), and perfused transcardially with 30 ml of PBS containing EDTA. Brains were removed and embedded in O.C.T. compound (Tissue-Tek) on dry ice and cryo-sectioned at a thickness of 20 µm. The sections were fixed with 4% PFA for 10 min, blocked with 5% normal donkey serum (Vector Labs) with Triton (0.05%) for 1 h, and incubated with primary antibodies for NeuN and SMI-312, or fibrinogen and DyLight 649-lectin (**Supp. Table 5**) diluted in blocking solution overnight at 4° C. Secondary antibodies (**Supp. Table 5**) were incubated 1 h at room temperature. Sections were cover slipped using fluorescent mounting medium (Dako) and imaged with a Nikon A1R HD inverted confocal microscope with Galvano scanner using high resolution optical sections that were captured with a 20x objective. Post-imaging analysis was performed using ImageJ (Fiji) software.

#### NeuN-positive neuronal nuclei counting

NeuN-positive neurons were imaged with a 20x objective in tissue sections adjacent to the injection site in the CA1 region of the hippocampus. Quantification of images were performed on a randomly selected region of interest (ROI) at 256×256 px (159.33×159.33 µm) by using the ImageJ Cell Counter analysis tool, as we previously described ^55^. The counted number of NeuN-positive cells was normalized per mm2 of brain tissue.

#### SMI-312-positive neuritic density analysis

SMI-312 signals were acquired, and ROI was selected following the same procedure as for NeuN images. SMI-312 signals were filtered using the median filter (radius =2), and manually thresholded to generate binary images by a blinded investigator using ImageJ (Fiji). The areas occupied by the signal in the binarized images were then quantified using the ImageJ Area measurement tool. The quantified areas in all images were normalized to account for background threshold adjustments. Total SMI-312-positive area was expressed as a percentage of total brain tissue area per each field, as we previously described ^55^.

#### Extravascular fibrinogen analysis

For detection of extravascular brain capillary fibrinogen deposits, an antibody that detects both fibrinogen and fibrinogen-derived fibrin polymers was used (**Supp. Table 5**). Quantification of images were performed on randomly selected 159.33×159.33 µm ROIs in somatosensory cortex and hippocampus. The fibrinogen-positive perivascular signal on the abluminal side of lectin-positive endothelial profiles on microvessels was analyzed by first manually thresholding images to generate binary images ImageJ (Fiji). The binarized areas occupied by the lectin-positive microvessel signals were subtracted from the binarized fibrinogen signal, and then the resulting fibrinogen signal was quantified using ImageJ Integrated density analysis measurement tool, similar to previously described ^23^.

### Clearance of amyloid and tau by iPSC-PC

#### Clearance of aggregated Aβ42 deposits

The Shandon multi-spot microscope glass slides (Thermo Scientific, 9991090) were coated with Cy3-labelled human Aβ42 at 5 µg per spot, as described previously ^32,33,60^. Spots were plated with 2,500 GFP-iPSC-PC per spot that were transduced with either with LRP1 siRNA (si.*LRP1*) (Horizon Discovery, E-004721-00-0020), LDLR siRNA (si.*LDLR*) (Horizon Discovery, E-011073-01-0020), VLDLR siRNA (si.*VLDLR*) (Horizon Discovery, 003721-00-0020), APOER2 siRNA (si.*APOER2*) (Horizon Discovery, E-011802-00-0020) or scrambled siRNA (si.*Control*) (Horizon Discovery, 001910-01-20), or left empty of cells. Cells were incubated for five days in pericyte culture medium (Sciencell Research Laboratories, 1201) and then fixed with 4% paraformaldehyde. Slides were scanned using a BZ-9000 fluorescence microscope with 20x objective and accompanying software. Three randomly chosen regions of interest within each spot from three independent cultures per group were analyzed, and the relative Cy3-Aβ42 signal intensity determined with the NIH ImageJ software.

#### Clearance of tau

GFP-iPSC-PC transduced with si*.Control* or si*.LRP1* were cultured on poly-L-lysine coated glass-bottom dishes and incubated with Cy3-labeled recombinant human tau protein (Cy3-tau) (R&D Systems, SP-495) in pericyte culture medium (Sciencell Research Laboratories, 1201) at 1 µg/ml for 48 h. The cell lysosomes were then labeled with LysoView 405 (Biotium Inc, 70066) in phenol red-free DMEM (Gibco 31053-028) for 30 min immediately before imaging. Z-stack images (10 mm height with 1 mm step size, 1024×1024 pixel resolution) were acquired at room temperature using a Nikon A1R HD inverted confocal microscope with 20x objective in Galvano scanning mode, and Nikon NIS Elements software. Five randomly chosen regions of interest (ROIs) within each dish from three independent cultures per group were analyzed. Max intensity projections of each Z-stack were generated in Fiji (ImageJ) software, thresholded, and area of Cy3-tau signal colocalized with LysoView-positive lysosomes was determined. The Cy3-positive-LysoView-positve area was normalized to the total area of GFP-positive iPSC-PC cells in the same image. Results from ROIs were averaged for each replicate.

#### Clerance of amyloid and tau (24h incubation)

For both Aβ42 and tau uptake assays, we followed the procedures described above for *Clearance of tau*, but evaluated at 24h instead of 48h. Briefly, iPSC-PC were cultured on poly-L-lysine coated glass-bottom dishes and labeled with CellTracker green or red. Cells were incubated with either FAM-Aβ42 or Cy3-labeled recombinant human tau protein (Cy3-tau) (R&D Systems, SP-495) in pericyte culture medium at 1 µg/ml for 24 h. The cell lysosomes were then labeled with LysoView 405 (Biotium Inc, 70066) in phenol red-free DMEM (Gibco 31053-028) for 30 min immediately before imaging. Z-stack images (10 mm height with 1 mm step size, 1024×1024 pixel resolution) were acquired at room temperature using a Nikon A1R HD inverted confocal microscope with 20x objective in Galvano scanning mode, and Nikon NIS Elements software. Images presented are max intensity projections of the Z-stacks acquired, and were generated in Fiji (ImageJ) software.

#### Amyloid uptake assay from brain sections

iPSC-PC were labeled with CellTracker Green CMFDA (Invitrogen, C7025) and cultured for at least 1 week before the brain section preparation. Cryosections (20 μm) from 10-month-old *5xFAD* mouse brains and APOE4-P301S were mounted on poly-L-lysine (PLL)-coated coverslips and stored at −80°C. On the day of the experiment, sections were thawed, rinsed with PBS, and incubated in pericyte culture medium (ScienCell) for 2 hours at 37°C. iPSC-PC were seeded onto tissue sections at 250,000 cells/well, with ‘no-cell’ controls included. Cultures were maintained for 24 hours at 37°C. Sections were fixed in 4% PFA, permeabilized in PBS containing donkey serum and 0.075% Triton X-100, and stained with a pan-Aβ antibody (1:1200, Cell Signaling Technology, 8243) or tau antibody (Phospho-Tau (Ser202, Thr205), 1:500 ThermoFisher, MN1020) followed by secondary antibody incubation (**Supp. Table 5**). Coverslips were mounted using DAPI-containing mounting medium and imaged using a confocal microscope at 4x and 10x magnification. For analysis, images were binarized using a customized threshold and percent of amyloid or tau signal were quantified in cortex and hippocampus using ImageJ (Fiji).

### Statistical Analysis

Data are presented as mean ± standard error of the mean (SEM), or standard deviation (SD) as indicated in figure captions. All analyses were performed using GraphPad Prism and RStudio 3.6.0. All data were tested for normal distribution using Shapiro-Wilk test. Two-tailed unpaired Student’s t-test was used to determine changes between two normally distributed groups. Statistical differences between multiple comparisons were determined using one-way analysis of variance (ANOVA) followed by the Tukey’s multiple comparisons test. Statistical significance was defined as ∗p < 0.05, ∗∗p < 0.01, and ∗∗∗p < 0.001. For further information regarding proteomic analysis please refer to the section above “*Statistics for proteomics analysis.*”

## Author Contributions

R.R., A.P.S., K.K., S.E.F., B.V.Z., and M.C. supervised the study and designed experiments. A.P.S., and C.G. generated iPSC-PC for experiments. A.P.S. performed western blot characterization and siRNA silencing of iPSC-PC. A.P.S. provided cells for proteomics studies. A.P.S., C.G. provided cells for functional in vitro and in vivo studies. J.TCW. contributed human iPSC lines. PC were sourced from the BioBank at the Center for Neurodegeneration and Regeneration at the Zilkha Neurogenetic Institute at USC. V.C. and M.C. prepared samples for LC/MS and performed all proteomics analyses. R.R. performed downstream proteomic analysis and visualization. A.P.S., K.K., and Y.K. performed *ex vivo* homing experiments and homing analysis on slices. Y.W. performed transplantation experiments. C.T.S. performed perfusions and brain tissue collection. R.R. and K.K. analyzed and visualized iPSC-PC homing in vivo. R.R. and A.S. performed high resolution imaging of grafted iPSC-PC and 3D reconstruction. R.R. and M.Z. analyzed brain tissue for BBB leakage and neuroprotection. A.P.S. and K.K. performed and analyzed Aβ and tau clearance studies in vitro. M.Z. and G.S. performed sectioning and staining of brain tissue. R.R. and M.Z. performed and analyzed Aβ and tau clearance on brain sections. R.R., A.P.S., and K.K. contributed to figure design and data visualization throughout the study. R.R., A.P.S., K.K., B.V.Z., and M.C. contributed to the writing of the manuscript, with input from all co-authors. All authors read and approved the final version of the manuscript.

## Acknowledgements and grants

This work is supported by the National Institutes of Health grant RF1AG039452, the Cure Alzheimer’s Fund award 012032-00001, and the Foundation Leducq Transatlantic Network of Excellence for the Study of Perivascular Spaces in Small Vessel Disease (reference no. 16 CVD 05). RR acknowledges funding support from Swiss 3R Competence Center (OC-2020-002), the Swiss National Science Foundation (CRSK-3_195902), (PZ00P3_216225), and Keck School of Medicine (KSOM) Dean’s Pilot Funding Program Award.

## References

1. Armulik, A. et al. Pericytes regulate the blood–brain barrier. Nature 468, 557–561 (2010).

2. Daneman, R., Zhou, L., Kebede, A. A. & Barres, B. A. Pericytes are required for blood–brain barrier integrity during embryogenesis. Nature 468, 562–566 (2010).

3. Bell, R. D. et al. Pericytes Control Key Neurovascular Functions and Neuronal Phenotype in the Adult Brain and during Brain Aging. Neuron 68, 409–427 (2010).

4. Nikolakopoulou, A. M. et al. Pericyte loss leads to circulatory failure and pleiotrophin depletion causing neuron loss. Nat Neurosci 22, 1089–1098 (2019).

5. Ayloo, S. et al. Pericyte-to-endothelial cell signaling via vitronectin-integrin regulates blood-CNS barrier. Neuron 110, 1641–1655.e6 (2022).

6. Sweeney, M. D., Zhao, Z., Montagne, A., Nelson, A. R. & Zlokovic, B. V. Blood-Brain Barrier: From Physiology to Disease and Back. Physiological Reviews 99, 21–78 (2019).

7. Tachibana, M., Yamazaki, Y., Liu, C.-C., Bu, G. & Kanekiyo, T. Pericyte implantation in the brain enhances cerebral blood flow and reduces amyloid-β pathology in amyloid model mice. Experimental Neurology 300, 13–21 (2018).

8. Ojo, J. et al. Mural cell dysfunction leads to altered cerebrovascular tau uptake following repetitive head trauma. Neurobiology of Disease 150, 105237 (2021).

9. Barisano, G. et al. A “multi-omics” analysis of blood–brain barrier and synaptic dysfunction in APOE4 mice. Journal of Experimental Medicine 219, e20221137 (2022).

10. Montagne, A. et al. APOE4 leads to blood–brain barrier dysfunction predicting cognitive decline. Nature 581, 71–76 (2020).

11. Faal, T. et al. Induction of Mesoderm and Neural Crest-Derived Pericytes from Human Pluripotent Stem Cells to Study Blood-Brain Barrier Interactions. Stem Cell Reports 12, 451–460 (2019).

12. Stebbins, M. J. et al. Human pluripotent stem cell–derived brain pericyte–like cells induce blood-brain barrier properties. Science Advances 5, eaau7375 (2019).

13. Sun, J. et al. Transplantation of hPSC-derived pericyte-like cells promotes functional recovery in ischemic stroke mice. Nat Commun 11, 5196 (2020).

14. Blanchard, J. W. et al. Reconstruction of the human blood–brain barrier in vitro reveals a pathogenic mechanism of APOE4 in pericytes. Nat Med 26, 952–963 (2020).

15. Winkler, E. A., Bell, R. D. & Zlokovic, B. V. Central nervous system pericytes in health and disease. Nat Neurosci 14, 1398–1405 (2011).

16. Li, J. et al. Long-term potentiation modulates synaptic phosphorylation networks and reshapes the structure of the postsynaptic interactome. Science Signaling 9, rs8–rs8 (2016).

17. Li, J. et al. Spatiotemporal profile of postsynaptic interactomes integrates components of complex brain disorders. Nat Neurosci 20, 1150–1161 (2017).

18. Wilkinson, B., Li, J. & Coba, M. P. Synaptic GAP and GEF Complexes Cluster Proteins Essential for GTP Signaling. Sci Rep 7, 5272 (2017).

19. Wilkinson, B. et al. Endogenous Cell Type–Specific Disrupted in Schizophrenia 1 Interactomes Reveal Protein Networks Associated With Neurodevelopmental Disorders. Biological Psychiatry 85, 305–316 (2019).

20. Kanev, G. K. et al. The Landscape of Atypical and Eukaryotic Protein Kinases. Trends in Pharmacological Sciences 40, 818–832 (2019).

21. Hornbeck, P. V. et al. PhosphoSitePlus: a comprehensive resource for investigating the structure and function of experimentally determined post-translational modifications in man and mouse. Nucleic Acids Res 40, D261–D270 (2012).

22. Gonzales, A. L. et al. Contractile pericytes determine the direction of blood flow at capillary junctions. Proc Natl Acad Sci U S A 117, 27022–27033 (2020).

23. Nikolakopoulou, A. M., Zhao, Z., Montagne, A. & Zlokovic, B. V. Regional early and progressive loss of brain pericytes but not vascular smooth muscle cells in adult mice with disrupted platelet-derived growth factor receptor-β signaling. PLOS ONE 12, e0176225 (2017).

24. Armulik, A., Genové, G. & Betsholtz, C. Pericytes: Developmental, Physiological, and Pathological Perspectives, Problems, and Promises. Developmental Cell 21, 193–215 (2011).

25. Smyth, L. C. D. et al. Characterisation of PDGF-BB:PDGFRβ signalling pathways in human brain pericytes: evidence of disruption in Alzheimer’s disease. Commun Biol 5, 1–16 (2022).

26. Zhang, H., Chen, H., Wang, W., Wei, Y. & Hu, S. Cell survival and redistribution after transplantation into damaged myocardium. J Cell Mol Med 14, 1078–1082 (2010).

27. Darsalia, V. et al. Cell number and timing of transplantation determine survival of human neural stem cell grafts in stroke-damaged rat brain. J Cereb Blood Flow Metab 31, 235–242 (2011).

28. Montagne, A. et al. Pericyte degeneration causes white matter dysfunction in the mouse central nervous system. Nature Medicine 24, 326–337 (2018).

29. Vanlandewijck, M. et al. A molecular atlas of cell types and zonation in the brain vasculature. Nature 554, 475–480 (2018).

30. Deane, R. et al. LRP/amyloid beta-peptide interaction mediates differential brain efflux of Abeta isoforms. Neuron 43, 333–344 (2004).

31. Zhao, Z. et al. Central role for PICALM in amyloid-β blood-brain barrier transcytosis and clearance. Nat Neurosci 18, 978–987 (2015).

32. Bell, R. D. et al. SRF and myocardin regulate LRP-mediated amyloid-beta clearance in brain vascular cells. Nat Cell Biol 11, 143–153 (2009).

33. Ma, Q. et al. Blood-brain barrier-associated pericytes internalize and clear aggregated amyloid-β42 by LRP1-dependent apolipoprotein E isoform-specific mechanism. Mol Neurodegener 13, 57 (2018).

34. Rauch, J. N. et al. LRP1 is a master regulator of tau uptake and spread. Nature 580, 381–385 (2020).

35. Oakley, H. et al. Intraneuronal β-Amyloid Aggregates, Neurodegeneration, and Neuron Loss in Transgenic Mice with Five Familial Alzheimer’s Disease Mutations: Potential Factors in Amyloid Plaque Formation. J. Neurosci. 26, 10129–10140 (2006).

36. Shi, Y. et al. ApoE4 markedly exacerbates tau-mediated neurodegeneration in a mouse model of tauopathy. Nature 549, 523–527 (2017).

37. Winkler, E. A. et al. A single-cell atlas of the normal and malformed human brain vasculature. Science 375, eabi7377 (2022).

38. Yang, A. C. et al. A human brain vascular atlas reveals diverse mediators of Alzheimer’s risk. Nature 603, 885–892 (2022).

39. Guo, M. et al. Single cell RNA analysis identifies cellular heterogeneity and adaptive responses of the lung at birth. Nat Commun 10, 37 (2019).

40. Oliveira, F., Bondareva, O., Rodríguez-Aguilera, J. R. & Sheikh, B. N. Cultured brain pericytes adopt an immature phenotype and require endothelial cells for expression of canonical markers and ECM genes. Front. Cell. Neurosci. 17, (2023).

41. Gaceb, A., Özen, I., Padel, T., Barbariga, M. & Paul, G. Pericytes secrete pro-regenerative molecules in response to platelet-derived growth factor-BB. J Cereb Blood Flow Metab 38, 45–57 (2018).

42. Tcw, J. et al. Cholesterol and matrisome pathways dysregulated in astrocytes and microglia. Cell 185, 2213–2233.e25 (2022).

43. Sagare, A. P., Sweeney, M. D., Makshanoff, J. & Zlokovic, B. V. Shedding of soluble platelet-derived growth factor receptor-β from human brain pericytes. Neuroscience Letters 607, 97–101 (2015).

44. Verbeek, M. M., Otte-Höller, I., Wesseling, P., Ruiter, D. J. & de Waal, R. M. Induction of alpha-smooth muscle actin expression in cultured human brain pericytes by transforming growth factor-beta 1. Am J Pathol 144, 372–382 (1994).

45. Wu, Z. et al. Role of the MEOX2 homeobox gene in neurovascular dysfunction in Alzheimer disease. Nat Med 11, 959–965 (2005).

46. Chow, N. et al. Serum response factor and myocardin mediate arterial hypercontractility and cerebral blood flow dysregulation in Alzheimer’s phenotype. Proceedings of the National Academy of Sciences 104, 823–828 (2007).

47. Davis, J., Wagner, M. R., Zhang, W., Xu, F. & Nostrand, W. E. V. Amyloid β-Protein Stimulates the Expression of Urokinase-type Plasminogen Activator (uPA) and Its Receptor (uPAR) in Human Cerebrovascular Smooth Muscle Cells *. Journal of Biological Chemistry 278, 19054–19061 (2003).

48. Otsu, N. A Threshold Selection Method from Gray-Level Histograms. IEEE Transactions on Systems, Man, and Cybernetics 9, 62–66 (1979).

49. Weber, R. Z., et al. Neural xenografts contribute to long-term recovery in stroke via molecular graft-host crosstalk. Nature Communications (in Press), (2025).

50. Weber, R. Z. et al. A molecular brain atlas reveals cellular shifts during the repair phase of stroke. J Neuroinflammation 22, 112 (2025).

51. Rust, R. Ischemic stroke-related gene expression profiles across species: a meta-analysis. J Inflamm 20, 21 (2023).

52. Tallquist, M. D., French, W. J. & Soriano, P. Additive effects of PDGF receptor beta signaling pathways in vascular smooth muscle cell development. PLoS Biol 1, E52 (2003).

53. Winkler, E. A., Bell, R. D. & Zlokovic, B. V. Pericyte-specific expression of PDGF beta receptor in mouse models with normal and deficient PDGF beta receptor signaling. Mol Neurodegener 5, 32 (2010).

54. Percie du Sert, N., et al. The ARRIVE guidelines 2.0: Updated guidelines for reporting animal research*. J Cereb Blood Flow Metab 40, 1769–1777 (2020).

55. Kisler, K. et al. Pericyte degeneration leads to neurovascular uncoupling and limits oxygen supply to brain. Nat Neurosci 20, 406–416 (2017).

56. Wang, Y. et al. 3K3A-activated protein C stimulates postischemic neuronal repair by human neural stem cells in mice. Nat Med 22, 1050–1055 (2016).

57. Kawata, M. et al. Long-range axonal projections of transplanted mouse embryonic stem cell-derived hypothalamic neurons into adult mouse brain. PLoS One 17, e0276694 (2022).

58. Yasuhara, T. et al. Transplantation of human neural stem cells exerts neuroprotection in a rat model of Parkinson’s disease. J Neurosci 26, 12497–12511 (2006).

59. Paisey, S. J. et al. Imaging of human stem cell-derived dopamine grafts correlates with behavioural recovery and reveals microstructural brain changes. Neurobiol Dis 209, 106910 (2025).

60. Wyss-Coray, T. et al. Adult mouse astrocytes degrade amyloid-β in vitro and in situ. Nat Med 9, 453–457 (2003).

