## Supplemental Figure 1-12 for "Molecular signature and functional properties of human pluripotent stem cell-derived brain pericytes"

### Supplementary Figures

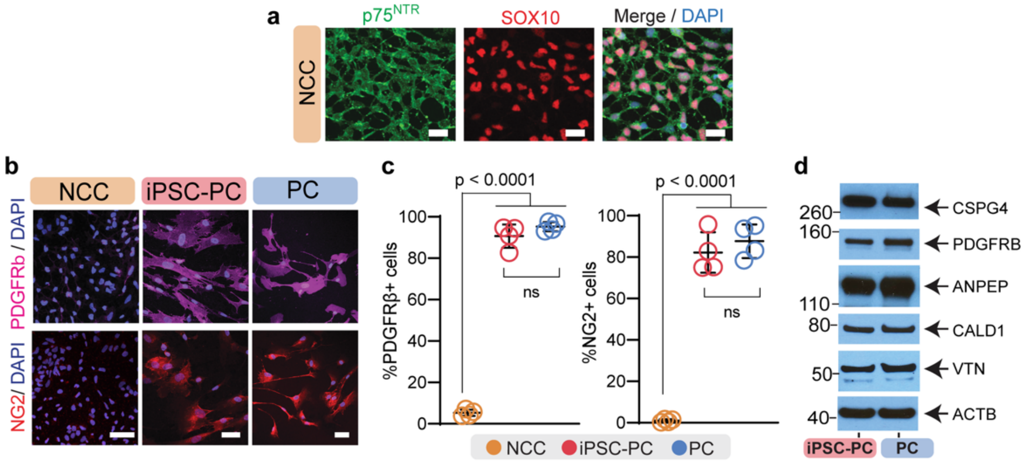


**Supplemental Figure 1. Initial characterization of iPSC-PC**. (**a**) iPSC-derived neural crest cells (NCC) express NCC markers p75 and SOX10. Scale bars = 20 μm. (**b-c**) Representative images (**b**) and quantification (**c**) of pericyte markers PDGFRb and NG2 in NCC, iPSC-PC and PC. In **b**, scale bars = 50 μm; DAPI, nuclear staining. In **c**, data presented as mean ± SEM, n = 4 independent cultures per group. Significance by Tukey's multiple comparisons test. (**d**) Immunoblotting of common pericyte markers NG2 (CSPG4), PDGFRβ (PDGFRB), CD13 (ANPEP), Caldesmon (CALD1), and Vitronectin (VTN) in iPSC-PC and PC. ACTB (b-actin) was used as a loading control.

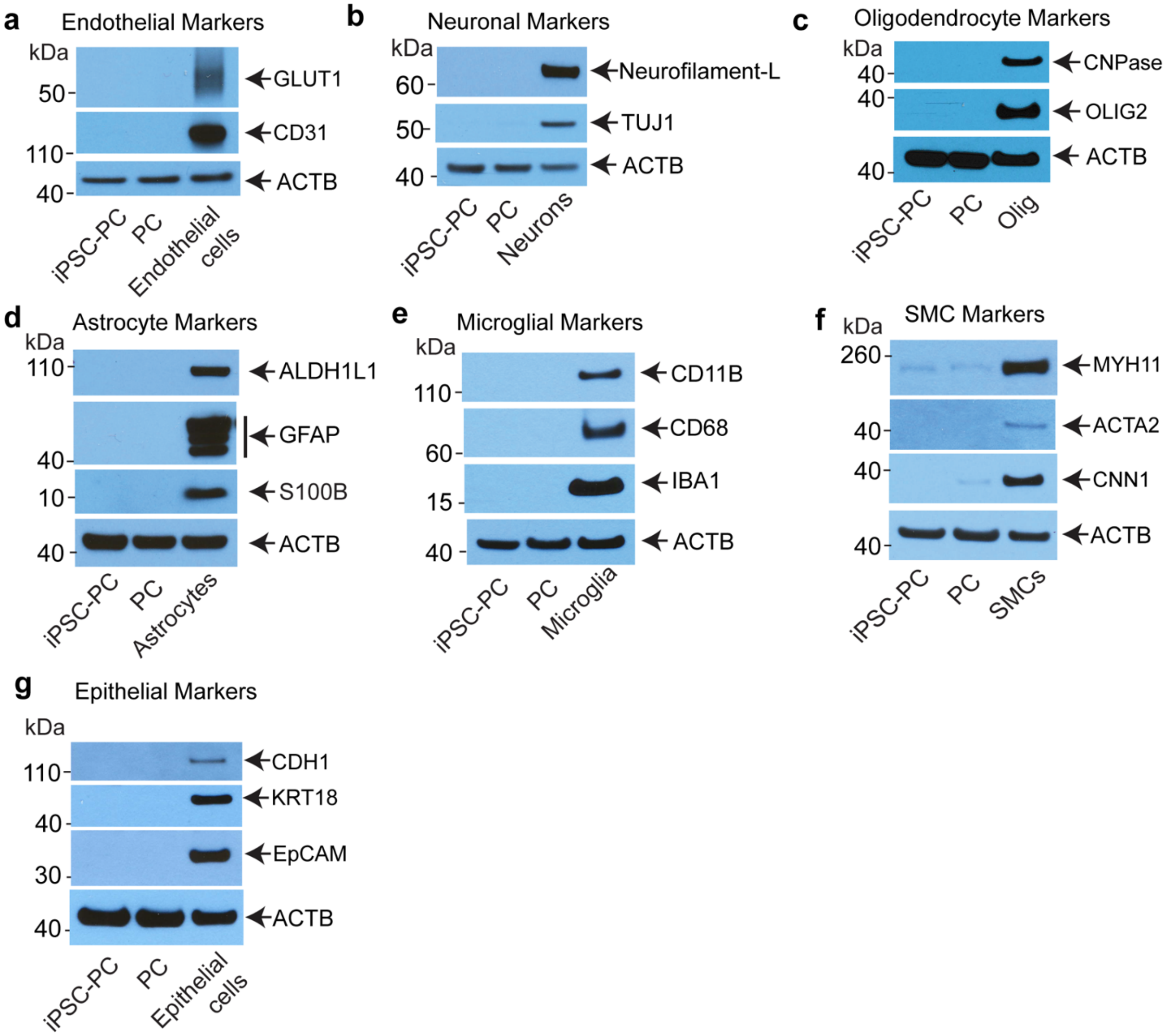


**Supplemental Figure 2. Evaluation of cell-specific CNS markers in iPSC-PC and PC.** Immunoblots illustrating that neither iPSC-PC nor PC express canonical markers of (**a**) endothelial cells, (**b**) neurons, (**c**) oligodendrocytes, (**d**) astrocytes, (**e**) microglia, or (**f**) SMC markers or **(g)** epithelial cell markers. ACTB (b-actin) was used as a loading control. iPSC-PC: induced pluripotent stem cell-derived pericytes, PC: primary human brain pericytes, SMC: smooth muscle cells, Olig: Oligodendrocytes, kDA: kilodalton.

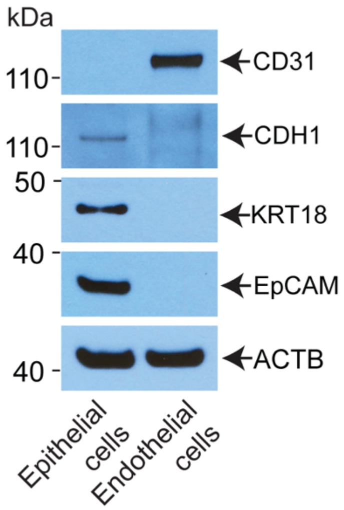


**Supplemental Figure 3: Evaluation of endothelial and epithelial specific markers in human primary cultured brain endothelial cells.** Immunoblots illustrating that primary human brain endothelial cells express the endothelial-specific marker CD31 and do not express epithelial cell markers (CDH1, KRT-19, EpCAM). Epithelial cells were used as a positive control. ACTB (b-actin) was used as a loading control.

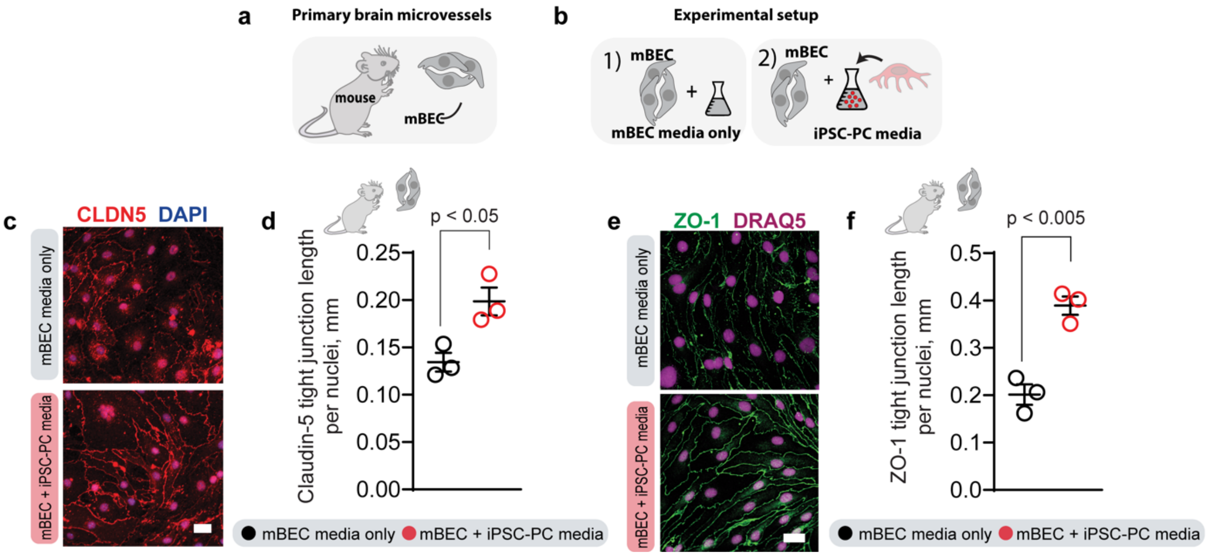


**Supplemental Figure 4. iPSC-PC extend the length of blood-brain barrier tight junction proteins.** (**a,b**) Schematic overview of experimental design using primary mouse brain endothelial cell (mBEC) cultures with different endothelial cell (mBEC) media formulations. (**c,d**) Representative images of tight junction Claudin-5 staining in mBEC cultures after 24 h treatment with iPSC-PC conditioned media or EC media only (**c**), and quantification of Claudin-5 tight junction protein length normalized to total number of DAPI nuclei (**d**). (**e,f**) Representative images of tight junction ZO-1 staining in mBEC cultures after 48 h treatment with iPSC-PC conditioned media or EC media only (**e**), and quantification of ZO-1 tight junction protein length normalized to total number of DRAQ5 nuclei (**f**). In **d**, **f**, data presented as mean ± SD; n = 3 replicates. Significance by unpaired t test. Scale bars: 25 μm.

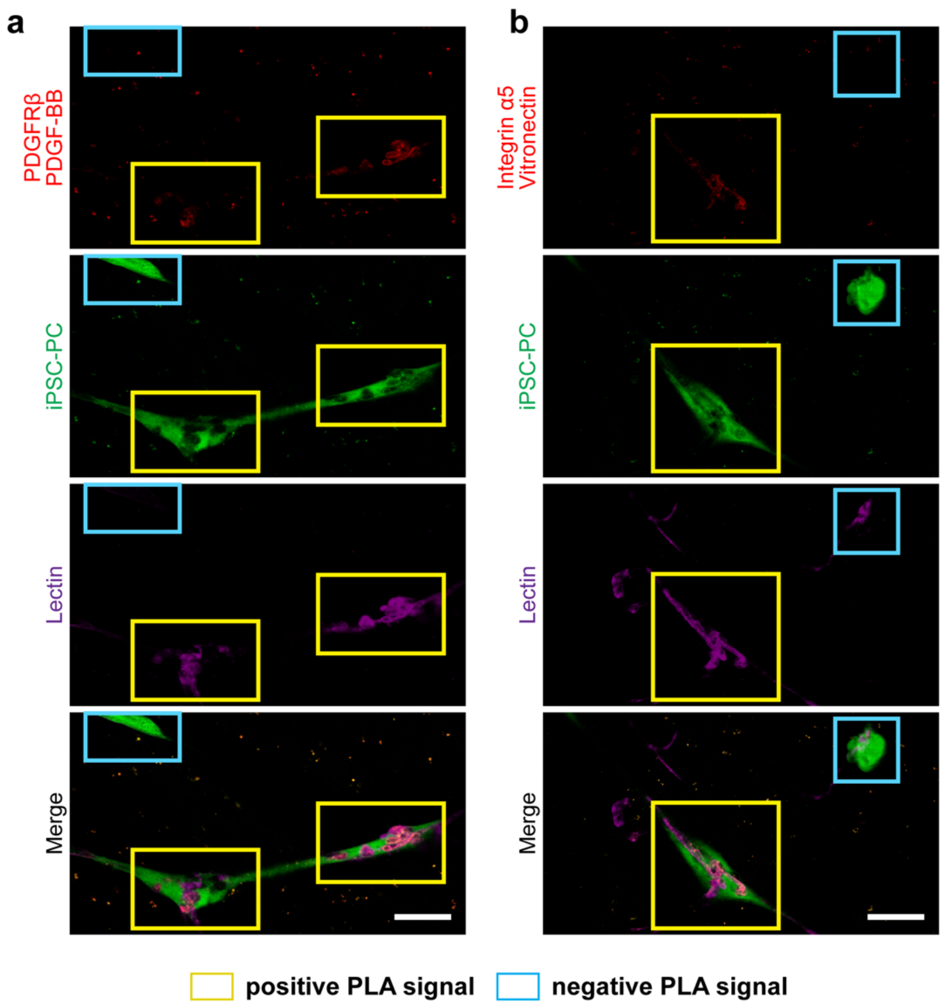


**Supplemental Fig. 5. Specificity of proximity ligation assay (PLA) for iPSC-PC–EC interactions.** Representative confocal images showing proximity labeling between iPSC-derived pericytes (iPSC-PC, green) and lectin+ endothelial cells (EC, magenta) for the ligand–receptor pairs PDGFRβ/PDGF-BB (left) and integrin α5/vitronectin (right). Yellow boxes highlight regions of direct iPSC-PC–EC contact showing PLA signal (red), whereas blue boxes mark non-associated iPSC-PCs lacking signal, confirming assay specificity. Small dispersed red puncta represent nonspecific background. Scale bars: 5 μm.

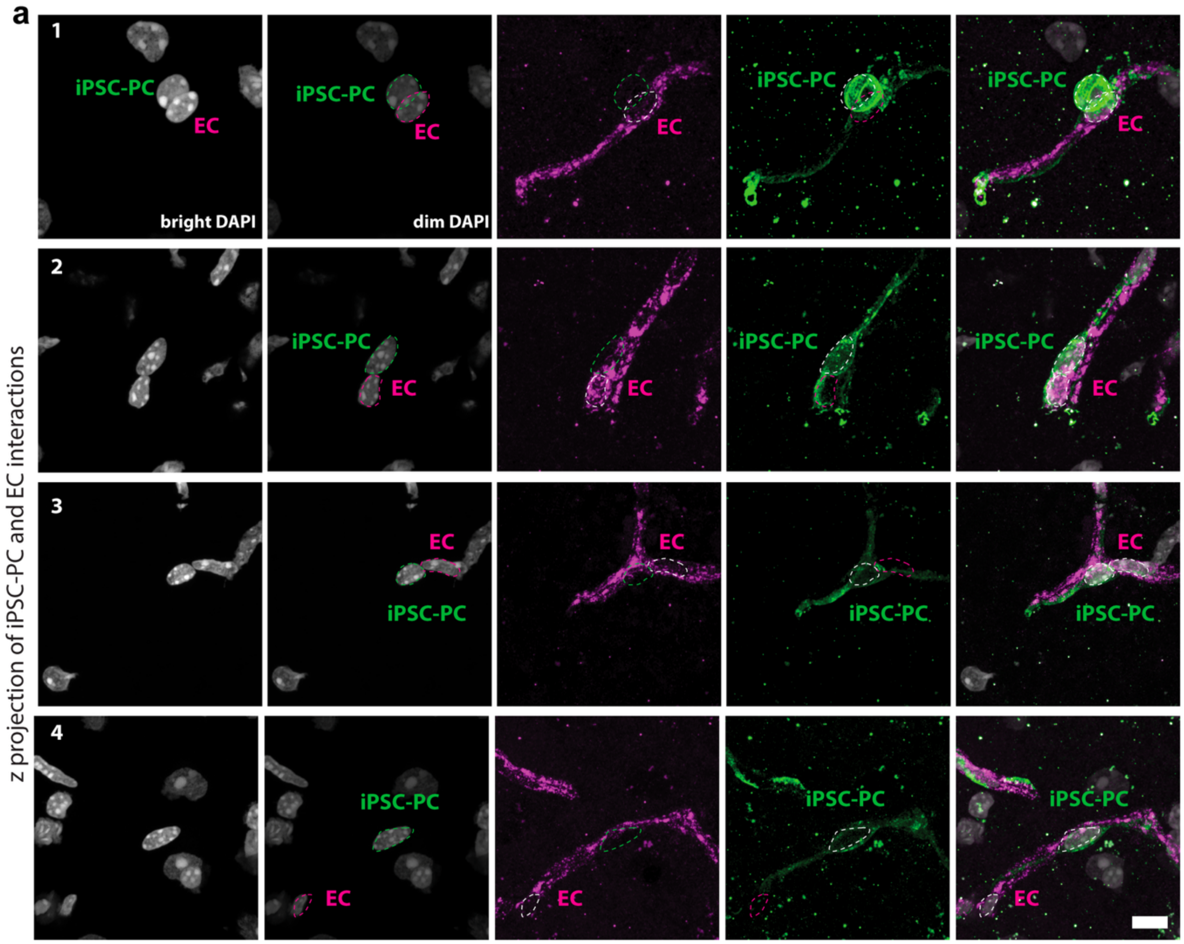


**Supplemental Figure 6. High-resolution z-projections of iPSC-PC and endothelial cell (EC) interactions 48 hours post-transplantation.** (a) Representative confocal z-projections showing iPSC-derived pericytes (iPSC-PC, green) interacting with endothelial cells (EC, magenta). Panels 1–4 show separate mice receiving iPSC-PCs with elongated morphologies with close proximity to ECs. Left to right: bright DAPI, dim DAPI, EC channel, iPSC-PC channel, and merged iPSC-PC/EC/DAPI images. Dashed circles indicate nuclei. Scale bar: 5 μm.

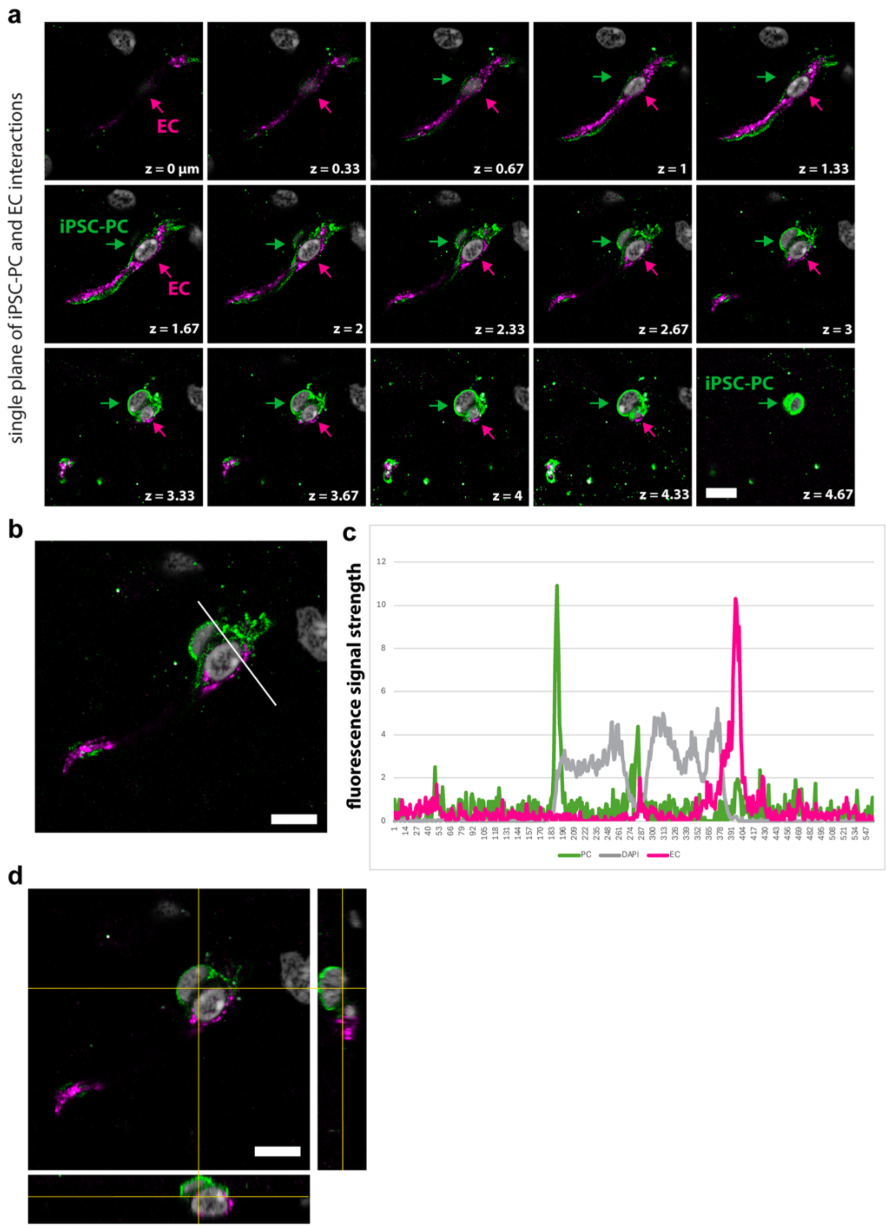


**Supplemental Fig. 7. Single-plane confocal z-stack of iPSC-PC and EC interactions.** Sequential optical sections (z = 0–4.67 µm) showing iPSC-derived pericytes (iPSC-PC, green) extending processes along endothelial cells (EC, magenta). Arrows indicate regions of nuclei across z-planes, highlighting gradual wrapping and alignment along the vessel surface. (b) Representative confocal image showing iPSC-derived pericytes (iPSC-PC, green) and endothelial cells (EC, magenta) at z = 3 µm. (c) Fluorescence intensity profile along the indicated white line in (b) demonstrating distinct spatial peaks for iPSC-PC (green) and EC (magenta) signals relative to nuclei (DAPI, gray). (d) Orthogonal view confirming spatial separation of iPSC-PC and EC signals across x–z and y–z planes. Scale bars: 5 µm.

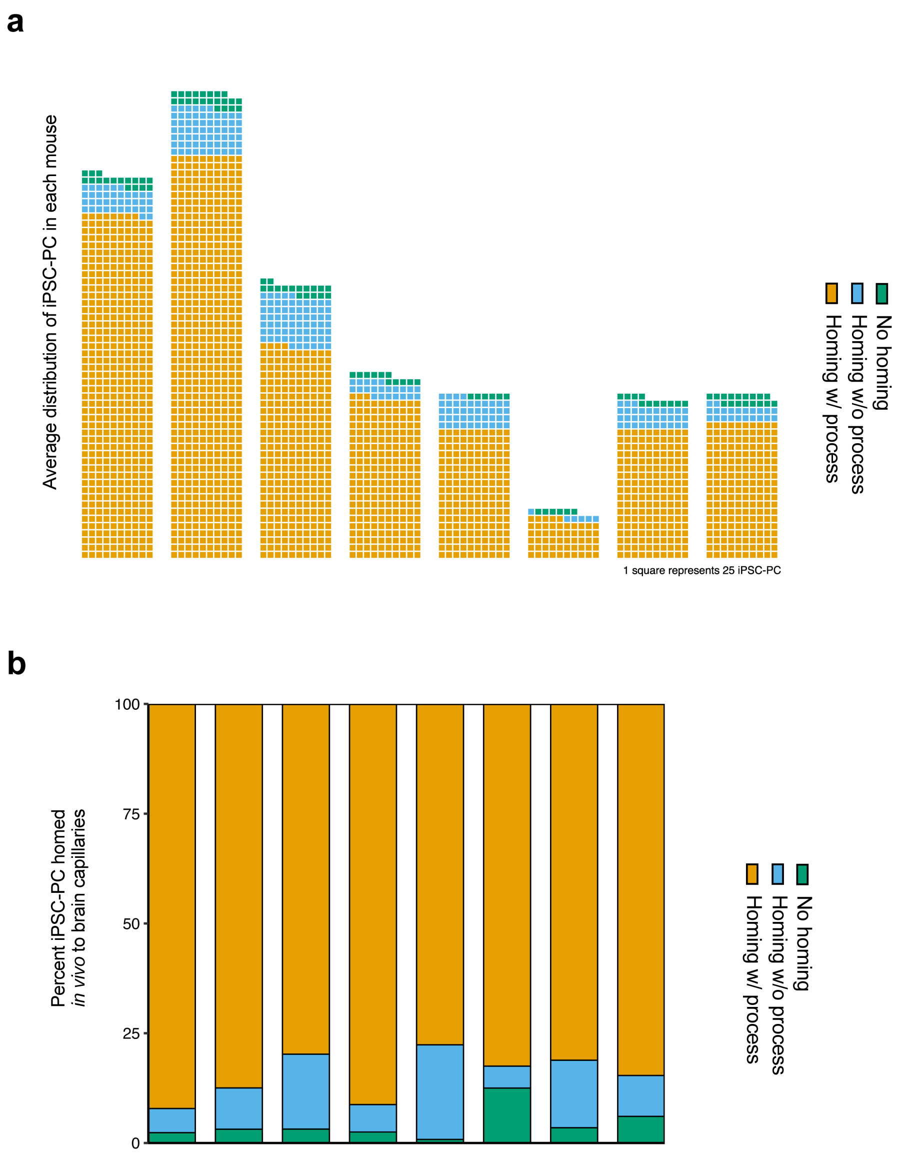


**Supplemental Figure 8: Quantification of iPSC-PC homing and vessel association in vivo.** (**a**) Estimated total number of surviving iPSC-PCs per animal based on counts across serial sections. Each square represents 25 iPSC-PC. Number above bar represents extrapolation of total count of iPSC-PC per mouse (**b**) Stacked bar showing proportions of iPSC-derived pericytes (iPSC-PC) classified as homing with processes (orange), homing without processes (blue), or not homing (green) to brain vessels 48 h post-transplantation Each bar is one animal in **a**,**b**.

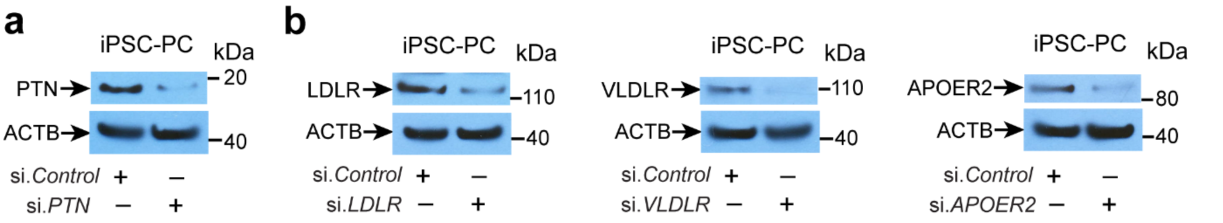


**Supplemental Figure 9. *PTN* silencing and lipoprotein receptor silencing in iPSC-PCs.** (**a**) Immunoblotting for PTN in iPSC-PC transduced with either control siRNA (si.*Control*) or human *PTN* siRNA (si.*PTN*). (**b**) Immunoblotting for lipoprotein receptors LDLR, VLDLR, APOER2 in iPSC-PC transduced with either si.*Control* or *si.LDLR, si.VLDLR, or si.APOER2*. For **a** and **b**, ACTB (β-actin) was used as a loading control.

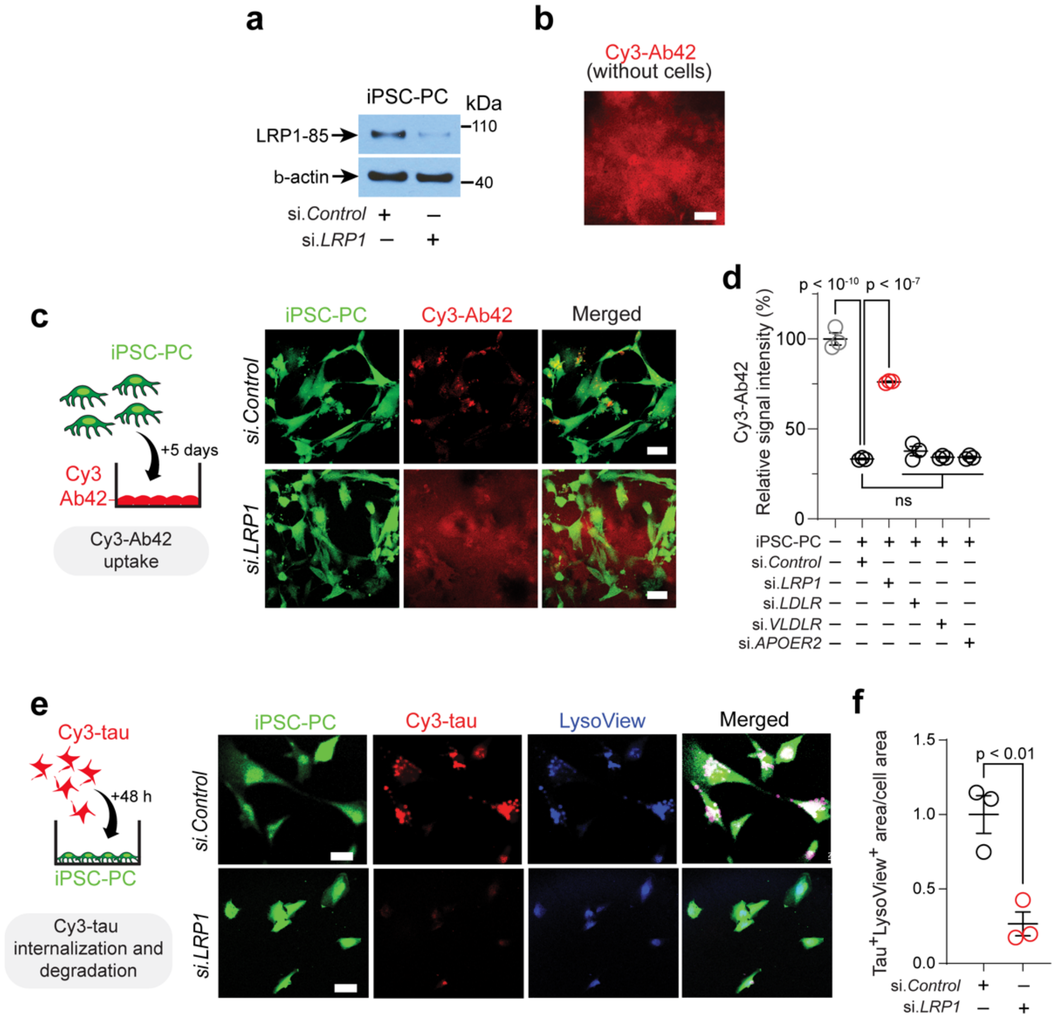


**Supplemental Figure 10. Aβ and tau clearance by iPSC-PC *in vitro*.** (**a**) Immunoblotting for *LRP1* in iPSC-PC transduced with either control siRNA (si.*Control*) or human *LRP1* siRNA (si.*LRP1*). (**b**) Representative confocal images of multi-spot glass slides coated with Cy3-labeled Aβ42 (red) without cells. (c) Experimental design (left) and representative confocal images (right) of iPSC-PC (green) after siRNA silencing of LRP1 (si.*LRP1*) or control siRNA (si.*Control*) added to Cy3-labeled Aβ42 coated multi-spot glass slides. (**d**) Quantification of Cy3-Aβ42 clearance by iPSC-PCs after silencing lipoprotein receptors *LRP1* (si.*LRP1*), *LDLR* (si.*LDLR*), *VLDLR* (si.*VLDLR*), and *APOER2* (si.*APOER2*) compared to si.*Control*. (**e,f**) Experimental design (left) and representative images (right) of human recombinant Cy3-tau (red) and LysoView 405-positive lysosomes (blue) after 48h incubation with iPSC-PC (green) that underwent siRNA silencing of *Lrp1* (si.*LRP1*) or control siRNA (si.*Control*) (**e**), and quantification of lysosomal tau colocalization (**f**). In **f**, data presented as fraction of $tau^+LysoView^+$ area relative to total cell area, with all data points normalized to the control mean. Data in **d**, **f** is mean ± SEM, with n = 3 cultures per condition. Significance by one-way ANOVA followed by Tukey's posthoc test in **d,** and two-tailed t-test in **f**. In **b**, **c**, **e**, scale bar = 25 μm.

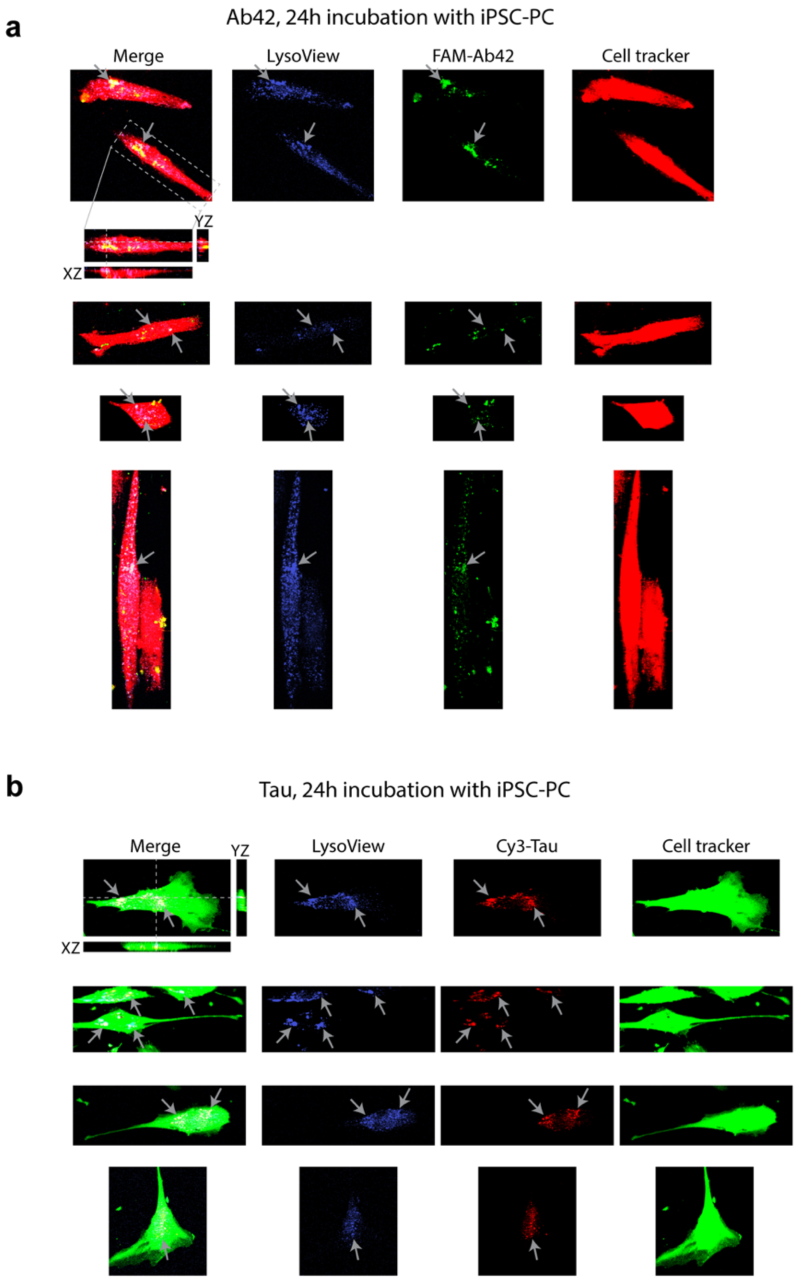


**Supplemental Fig. 11: Amyloid and tau clearance by iPSC-PC in vitro after 24h incubation *in vitro*.** (a) Representative images of human recombinant FAM-Aβ42 (green) and LysoView 405-positive lysosomes (blue) after 24h incubation with CellTracker red-labeled iPSC-PC (red). Gray arrows point out some locations were Aβ42 colocalizes with LysoView-positive lysosomes inside the cells (white color indicates colocalization). Top example also shows orthogonal views of a cell, showing Aβ42 was internalized. Orthogonal views were enlarged 2x in the z-direction for clarity. Images are representative of 3 replicates. (b) Representative images of human recombinant Cy3-tau (red) and LysoView 405-positive lysosomes (blue) after 24h incubation with CellTracker green-labeled iPSC-PC (green). Gray arrows point out some locations were tau colocalizes with LysoView-positive lysosomes inside the cells (white color indicates colocalization). Top example also shows orthogonal views of a cell, showing tau was internalized. Orthogonal views were enlarged 2x in the z-direction for clarity. Images are representative of 4 replicates.

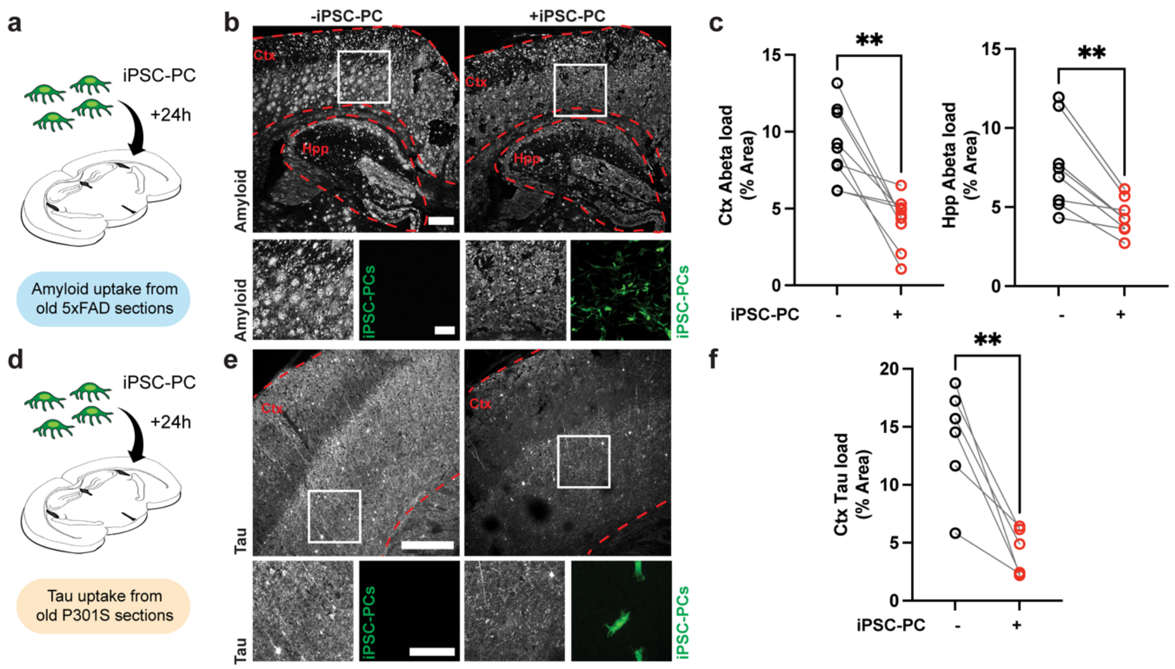


**Supplemental Figure 12: Clearance of Aβ and tau by iPSC-PC in 5xFAD and APOE4 P301S mouse brain sections.** (**a**) Schematic of amyloid uptake by iPSC-PC from 10 month old 5xFAD mouse brain sections. (**b-c**) Representative confocal images showing amyloid (white) in 5xFAD tissue sections after 24 h incubation with iPSC-PC (green) and control sections incubated in media without iPSC-PC (**b**), and quantification of cortical and hippocampal amyloid load in iPSC-PC treated sections and controls (**c**). In (**b**), boxes show locations of insets. (**d**) Schematic of tau uptake by iPSC-PC from 10 month old APOE4-P301S mouse brain sections. (**e-f**) Representative confocal images showing tau (white) in APOE4 P301S tissue sections after 24 h incubation with iPSC-PC (green) and control sections incubated in media without iPSC-PC (**e**), and quantification of hippocampal tau load in iPSC-PC treated sections and controls (**f**). In (**e**), boxes show locations of insets. Data in **c**, **f** represent paired consecutive sections from the same mouse brain, with pairs connected by lines; n = 6-7 sections. Each circle represents an individual section. Significance was assessed by a paired two-tailed t-test. Scale bars: 200 µm (**b,e** overview), 50 µm (**b,e**, insets).

### Supplementary Tables

**Supplemental Table 1: Proteomics raw data and metadata of iPSC-PC and PC.**

**Supplemental Table 2: Phosphopeptides raw data and metadata of iPSC-PC and PC.**

**Supplemental Table 3: Processed proteomics data and summary in iPSC-PC and PC.**

**Supplemental Table 4: Processed phosphopeptides data and summary in iPSC-PC and PC.**

### Supplementary Videos

**Supplemental Video 1:** 3D rendering of iPSC-PC transduced with *si.Control* (green) integrating with mouse brain capillaries (magenta) after 24 h incubation with brain slices.

**Supplemental Video 2:** 3D rendering of iPSC-PC transduced with *si.PDGFRB* (green) that do not associate with mouse brain capillaries (magenta) after 24 h incubation with brain slices.

**Supplemental Video 3:** 3D rendering of iPSC-PC (green) homing to host mouse vessels (magenta) 24 h post-transplantation from representative mouse 1.

**Supplemental Video 4:** 3D rendering of iPSC-PC (green) homing to host mouse vessels (magenta) 48 h post-transplantation from representative mouse 2.

**Supplemental Video 5:** 3D rendering of iPSC-PC (green) homing to host mouse vessels (magenta) 48 h post-transplantation from representative mouse 3.
